# Detecting macroevolutionary genotype-phenotype associations using error-corrected rates of protein convergence

**DOI:** 10.1101/2022.04.06.487346

**Authors:** Kenji Fukushima, David D. Pollock

## Abstract

On macroevolutionary timescales, extensive mutations and phylogenetic uncertainty mask the signals of genotype-phenotype associations underlying convergent evolution. To overcome this problem, we extended the widely used framework of nonsynonymous-to-synonymous substitution rate ratios and developed the novel metric *ω_C_*, which measures the error-corrected convergence rate of protein evolution. While *ω_C_* distinguishes natural selection from genetic noise and phylogenetic errors in simulation and real examples, its accuracy allows an exploratory genome-wide search of adaptive molecular convergence without phenotypic hypothesis or candidate genes. Using gene expression data, we explored over 20 million branch combinations in vertebrate genes and identified the joint convergence of expression patterns and protein sequences with amino acid substitutions in functionally important sites, providing hypotheses on undiscovered phenotypes. We further extended our method with a heuristic algorithm to detect highly repetitive convergence among computationally nontrivial higher-order phylogenetic combinations. Our approach allows bidirectional searches for genotype-phenotype associations, even in lineages that diverged for hundreds of millions of years.

## Introduction

A central aim of modern biology is to differentiate the huge amount of nonfunctional genetic noise from phenotypically important changes. Evolutionary processes at the molecular level are largely neutral and stochastic, but natural selection can constrain evolutionary pathways available to the organism. If similar environmental conditions recur in divergent lineages, the adaptive response may also be similar, leading to convergence, the repeated emergence of similar features in distantly related organisms (Losos, 2017). The prevalence of phenotypic convergence is demonstrated by various examples throughout the tree of life, such as the camera eyes of vertebrates and cephalopods, powered flight of birds and bats, and trap leaves of distantly related carnivorous plants. Because the repeated emergence of such complex traits by neutral evolution alone is extremely unlikely, convergence at the phenotypic level is considered strong evidence for natural selection.

Phenotypic convergence is necessarily caused by molecular events and often coincides with detectably excess levels of convergent molecular changes in gene regulation, gene sequences, gene repertoires, and other hierarchies of biological organization (Stern, 2013; Storz, 2016). A meta-analysis reported that 111 out of 1,008 loci had been convergently modified to attain common phenotypic innovations, sometimes even between different phyla (Martin and Orgogozo, 2013), demonstrating that genotype-phenotype associations frequently occur on macroevolutionary scales. For example, several lineages of mammals, reptiles, amphibians, and insects acquired resistance to toxic cardiac glycosides using largely overlapping sets of amino acid substitutions in a sodium channel (Ujvari et al., 2015). Another example illustrated how human cancer cells and plants employed common amino acid substitutions in Topoisomerase I to cope with a common toxic cellular environment generated by plant-derived anticancer drugs (Sirikantaramas et al., 2008).

Genome sequences are becoming more available for diverse lineages from the entire tree of life (Lewin et al., 2022), making it possible to explore macroevolutionary genotype-phenotype associations on large scales. However, because most molecular changes are nearly neutral or essentially nonfunctional in nature (Ohta, 1973), false-positive convergence in the form of stochastic, nonadaptive, convergent events is particularly problematic when conducting a genome-scale search. Furthermore, false positives can arise from methodological biases. For molecular convergence, a major source of bias occurs because such inference is sensitive to the topology of the phylogenetic tree on which substitution events are placed (Mendes et al., 2016) (Fig. 1A), while alternative methods that do not place substitutions on phylogenetic trees suffer even more severe rates of false positives (Foote et al., 2015; Thomas and Hahn, 2015; Zou and Zhang, 2015b) (Supplementary Text 1). A correctly inferred tree avoids false positives due to phylogeny (Castoe et al., 2009), but topological misinference due to technical errors, insufficient data, or biological factors such as introgression, horizontal gene transfer (HGT), paralogy, incomplete lineage sorting, and within-locus recombination, can all create substantial amounts of false convergence signals even when adaptive convergence did not actually occur (Mendes et al., 2016, 2019; Stern, 2013; Thomas et al., 2017). Importantly, false convergence events driven by topological errors tend to similarly affect both nonsynonymous and synonymous substitutions (Fig. S1A). By contrast, truly adaptive convergence should occur almost exclusively in nonsynonymous substitutions (amino acid–changing substitutions), as positive selection on synonymous substitutions is negligible, or at least not prevalent (Yang, 2006) (Fig. S1B). Therefore, synonymous convergence can potentially serve as a reliable reference for measuring the rate of expected nonsynonymous convergence due to phylogenetic inference error.

**Figure 1.**
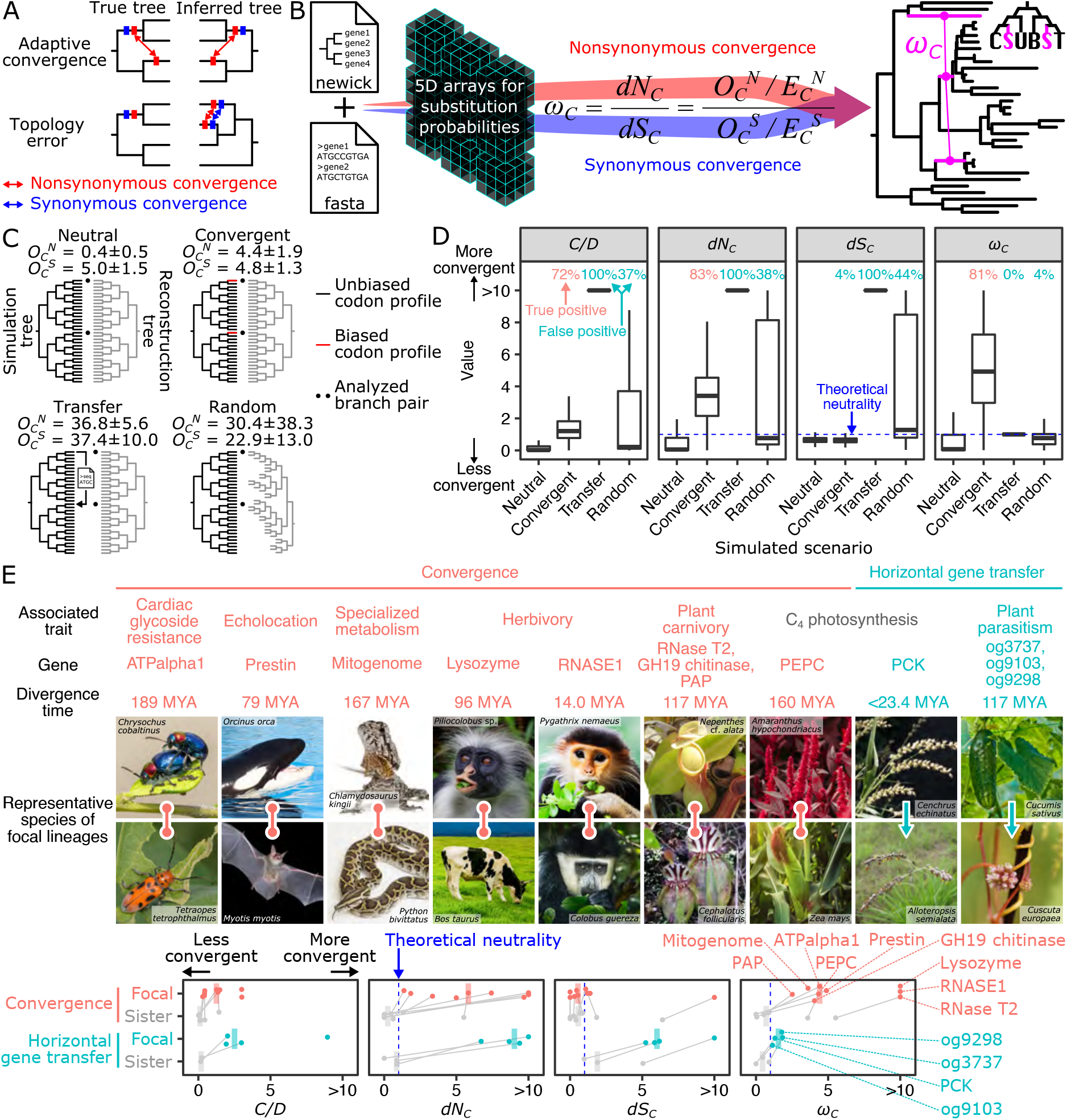
Challenges and solutions for the detection of molecular convergence. (**A**) False convergence is caused by tree topology errors. (**B**) The overview of CSUBST. This program processes substitution probabilities to derive observed (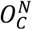 and 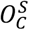) and expected (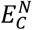 and 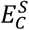) numbers of nonsynonymous and synonymous convergence and evaluate their rates (*dN_C_* and *dS_C_*) in branch combinations in a phylogenetic tree. A more detailed illustration is available in Fig. S2. (**C**) Generation of simulated datasets for performance evaluation in different evolutionary scenarios. The ECMK07+F codon substitution model was used to simulate the evolution of 500-codon sequences on a phylogenetic tree with 32 leaves 1,000 times. The numbers of observed nonsynonymous and synonymous convergence are indicated above trees (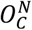 and 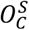, respectively; mean ± standard deviation). (**D**) The estimated rates of protein convergence in different scenarios. Each box plot corresponds to the results of 1,000 simulations. Dashed lines indicate the neutral expectation (=1.0) except for *C/D* (Castoe et al., 2009; Goldstein et al., 2015), for which no theoretical expectation is available. *dN_C_* is largely equivalent to the previously proposed metric called *R* (Zou and Zhang, 2015a). Values greater than the 95th percentile in the Neutral scenario are defined as true and false positives in Convergent and other scenarios, respectively, and are indicated at the top of the plot. The positive rate of *dS_C_* is interpreted as a false positive rate even in the Convergent scenario because the probability of only nonsynonymous substitutions is manipulated. (**E**) Performance of convergence metrics in empirical datasets. Known examples of protein convergences and horizontal gene transfers (HGTs) are analyzed with *C/D, dN_C_*, *dS_C_*, and *ω_C_*. Median values (bars) are overlaid on individual data points that correspond to gene trees. In trees where convergence occurred in more than two lineages, the median of all foreground branch pairs is reported. The branch pairs sister to the focal branches are shown as a control (Foote et al., 2015), except in cases where there is no substitution at all or the sister branches are phylogenetically not independent. Dataset and photographs of representative species are shown above the plot. The taxonomic range follows the NCBI Taxonomy database (Schoch et al., 2020), and the divergence time is according to timetree.org (Hedges et al., 2015). The lineages involving adaptive convergence or HGTs are referred to as focal lineages. The gene trees are illustrated in Fig. S5 and Fig. S6. The comparison with the background levels for each dataset is shown in Fig. S4. The characteristics of the datasets are summarized in Table S3. The photograph of *Alloteropsis semialata* is licensed under CC BY-SA 3.0 (https://creativecommons.org/licenses/by-sa/3.0/) by Marjorie Lundgren.

A widely used framework to understand how functionally constrained proteins evolve compared to neutral expectations is to contrast rates of nonsynonymous and synonymous substitutions. The ratio of these rates within a protein-coding sequence accounts for mutation biases and is often denoted as *ω*, *dN/dS*, or *K_a_/K_S_* (Zhang and Yang, 2015). Here, we extend this framework to derive the new metric ratio *ω_C_* and implement it to measure phylogenetic error-corrected rates of convergence. Simulation and empirical data analysis show that this new metric has high sensitivity while suppressing false positives. We further show its capability to detect factors that affect protein convergence rates and to identify likely adaptive protein evolution in a genome-scale dataset by an exploratory analysis without a pre-existing hypothesis. We also develop a heuristic algorithm to explore convergent signals with high signal-to-noise ratios in exponentially increasing numbers of higher-order phylogenetic combinations.

## Results

### Extending the framework of nonsynonymous per synonymous substitution rate ratio to molecular convergence

One of the most commonly accepted measures of the rate of protein evolution compared to neutral expectations is the ratio between nonsynonymous and synonymous substitution rates, denoted as *dN* and *dS*, or *K_a_* and *K_s_*, respectively (Yang, 2006). Using the ratio *dN/dS* to measure relative rates of protein evolution is justified, as the selective pressure on synonymous sites is negligible compared to that on nonsynonymous sites and thus remains fairly constant relative to the mutation rate (Yang, 2006). In a model-based framework, this ratio is parameterized as *ω*.

Inspired by *ω*, we developed a similar metric, *ω_C_*, that applies to substitutions that occurred repeatedly on a combination of separate phylogenetic branches (combinatorial substitutions; Fig. S1C; Supplementary Text 2). The metric *ω_C_* estimates the relative rates of convergence obtained by contrasting the rates of nonsynonymous and synonymous convergence (*dN_C_* and *dS_C_*, respectively). Using this ratio, important biological fluctuations, such as among-site rate heterogeneity and codon equilibrium frequencies, are taken into account (for details, see Supplementary Text 3 and Methods). Similar to previously proposed convergence metrics (Castoe et al., 2009; Goldstein et al., 2015; Zou and Zhang, 2015a), *ω_C_* is calculated from substitutions at multiple codon sites across protein-coding sequences. As a result, one *ω_C_* value is obtained for each gene for each branch pair (or for a combination of more than two branches) in the phylogenetic tree. A unique feature of *ω_C_* setting it apart from other metrics is its error tolerance. For example, if one of the branches in a branch combination is in error, *ω_C_* is a measure of the ratio of false convergence events of both kinds falsely attributed to a non-existent branch combination. In this way, the *ω_C_* values remain close to the neutral expectation of 1.0, even when topology errors are involved. Our method is implemented in the python program CSUBST (https://github.com/kfuku52/csubst), which takes as input a rooted phylogenetic tree and a codon sequence alignment (Fig. 1B and Fig. S2).

### The robustness of *ω_C_* as a relative rate of molecular convergence

Conventionally, observed levels of convergent amino acid substitutions have been contrasted either to the amount of convergence expected under a neutral model with no constraint (*R* (Zou and Zhang, 2015a)) or to other combinations of amino acid substitution patterns that are similarly affected by site-specific constraint (i.e., double divergence; *C/D* (Castoe et al., 2009; Goldstein et al., 2015)) (Table S1; Supplementary Text 4). The metric *R*, for example, is intended to have an expectation of 1.0 under neutral evolution, but in practice is somewhat lower than 1.0, even when the tree and substitution model are correct and exactly match simulation conditions (Zou and Zhang, 2015a). Using *R >* 1.0 as a criterion to identify convergence is thus in principle conservative for detecting convergence levels greater than fully neutral evolution. Furthermore, its accuracy depends on the accuracy of the phylogenetic tree in various aspects, e.g., neutral substitution model, tree topology, branch lengths, and reconstructed ancestral states. By contrast, the *C/D* comparison ratio, which compares convergence levels to double divergence events between branch pairs, is not strongly dependent on neutral substitution estimates (Castoe et al., 2009; Goldstein et al., 2015); however, it is dependent on the accuracy of the reconstructed tree compared to the true tree that applies. The *C/D* ratio may vary among proteins due to varying levels of constraint among proteins but is generally well below 1.0 (Goldstein et al., 2015).

Here, we focus on whether *ω_C_* performs better as a measure of convergence between branches in comparison to alternative metrics. Accordingly, we generated simulated sequences with 500 codons along a balanced phylogenetic tree ending with 32 sequences at the tips (or leaves), in all cases comparing two deeply separated tip lineages (shown as dots in Fig. 1C; Table S2). In this analysis, we compared *C/D, dN_C_*, *dS_C_*, and *ω_C_* under four evolutionary scenarios of relationships between the two tips being compared: 1) full neutral evolution along all branches (Neutral); 2) neutral evolution for nearly all branches but with convergent selection along the two deeply separated tip lineages (Convergent); 3) neutral evolution with phylogenetic tree topology error in the form of a copy-and-paste transfer from one of the two deeply separated lineages to the other, overwriting its genetic information (Transfer); or 4) neutral evolution but using a randomly reconstructed phylogenetic tree to detect convergence (Random). The metric *dN_C_* is obtained by dividing the observed value of nonsynonymous convergence 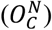 by the expected value 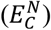 and is essentially equivalent to the previously proposed metric called *R* (Zou and Zhang, 2015a), but we use the *dN_C_* notation here to clarify its relationship to *dS_C_*, the ratio of observed to expected values of synonymous convergence 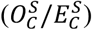.

During neutral evolution, sequences evolved under a constant codon substitution model without any adaptive convergence or constraint on amino acid substitutions other than those imposed by the structure of the genetic code and relative codon frequencies. In the Neutral scenario (Fig. 1C), the trees used for simulation and reconstruction were identical. *C/D* was much lower than 1.0, as expected, while the other three metrics (*dN_C_*, *dS_C_*, and *ω_C_*) were close to but lower than the neutral expectation of 1.0 (Fig. 1D). This observation is likely due to the fact that the convergent events must be inferred and are not actually observed, as investigated previously in *R* (Zou and Zhang, 2015a). In the Convergent scenario, adaptive convergence on the focal pair of deeply separated branches (red branches in Fig. 1C) was mimicked by convergently evolving 5% of codon sites (25 sites) in the two branches under substitution models biased toward codons encoding the same randomly selected amino acid. This generated an average of four excess nonsynonymous convergent substitutions on these two branch pairs (see 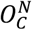 in Fig. 1C). In the Convergent scenario, the three protein convergence metrics, *C/D, dN_C_*, and *ω_C_*, yielded values substantially higher than they did under the Neutral scenario, while the synonymous change measure *dS_C_* remained comfortably well below 1.0. Using the distribution of metric values under the Neutral scenario as a reference, we see that 70–80% of the detection metric values in the Convergent scenario are above the 95^th^ percentile of the 1,000 simulations in their respective neutral distributions, while only 3.5% of *dS_C_* values are above this threshold, indicating that this level of convergence is usually detected by all three of the protein convergence metrics (Fig. 1D). To be thorough, we considered that *ω_C_* metrics can in general be derived for nine types of combinatorial substitutions (i.e., substitutions occurring at the same protein site in multiple independent branches; Supplementary Text 2) based on whether the ancestral and descendant states are the same or different, or in any state among multiple branches (Fig. S1C). In the Convergent scenario, only the *ω_C_* metrics involved in convergence (i.e., not divergence) showed a response, confirming its specificity (Fig. S3A).

We next considered Transfer and Random scenarios that include phylogenetic error. In the Transfer scenario, we transferred one of the focal tip sequences to the other focal tip sequence in the simulation, but the phylogenetic tree used in the analysis remained unchanged, as might happen with HGT events (Fig. 1C). In the Random scenario, we fully randomized the entire reconstructed tree relative to the true tree (Fig. 1C). Excess convergence detected in either of these scenarios is considered a false positive. We determined that both *C/D* and *dN_C_* are sensitive to the errors (Fig. 1D). By contrast, and as intended, *ω_C_* values were close to the neutral expectation because the rise in *dN_C_* due to phylogenetic error is matched by a similar increase of *dS_C_*, and they cancel each other out in the *ω_C_* metric (Fig. 1D). Further simulations supported the robustness of *ω_C_* against the rate of protein evolution, model misspecification, tree size, and protein size (Fig. S3B–F; Supplementary Text 5). Furthermore, *ω_C_* showed low false-positive rates in sister branches that serve as a control for the focal branch pairs (Foote et al., 2015) (Fig. S3G). Taken together, our simulation showed that *ω_C_* effectively counteracts false positives caused by phylogenetic errors without loss of power.

### *ω_C_* distinguishes between adaptive and false convergence in empirical datasets

To test whether *ω_C_* performs well with real data, we collected protein-coding sequence datasets from known molecular convergence events in various pairs of lineages covering insects, tetrapods, and flowering plants (Fig. 1E, Fig. S4, Fig. S5, and Table S3). Insects that feed on milkweed (Apocynaceae) harbor amino acid substitutions in a sodium pump subunit (ATPalpha1) that confer cardiac glycoside resistance (Dobler et al., 2012; Yang et al., 2019a; Zhen et al., 2012) (Fig. S4A). Echolocating bats and whales share amino acid substitutions in the hearing-related motor protein Prestin to enable high-frequency hearing (Liu et al., 2010, 2014) (Fig. S4B). An extensive molecular convergence occurred in the mitochondrial genomes of agamid lizards and snakes, presumably due to physiological adaptations for radical fluctuations in their aerobic metabolic rates (Castoe et al., 2009). Specialized digestive physiology of herbivorous mammals and carnivorous plants led to the molecular convergence of digestive enzymes (Fukushima et al., 2017; Stewart et al., 1987; Zhang, 2006; Zhang and Kumar, 1997) (Fig. S4C–G). Phosphoenolpyruvate carboxylase (PEPC), a key enzyme for carbon fixation in C_4_ photosynthesis, shares multiple amino acid convergence (Besnard et al., 2009; Christin et al., 2007) (Fig. S4H). In all these examples, *ω_C_* successfully detected convergent lineages, while it was always lower and in many cases close to the neutral expectation in the branch pairs sister to the focal lineages, which serve as a negative control (Fig. 1E; Table S4). Moreover, the *ω_C_* values of the focal branch pairs tended to be high compared to background levels in the phylogenetic trees (Fig. S4I). Analysis of different categories of combinatorial substitutions correctly recovered a trend consistent with the action of intramolecular epistasis, which did not appear in the simulations (Supplementary Text 6; Fig. S4J–K).

To test robustness against phylogenetic errors, we also employed reported cases of HGTs associated with C_4_ photosynthesis (Dunning et al., 2019) and plant parasitism (Yang et al., 2019b). We reconstructed the phylogenetic trees of the HGT genes with a constraint that enforces species tree-like topologies (Fig. S6). This operation separates the HGT donor and acceptor lineages and creates false convergence (Fig. S1A). Consistent with the simulation results, *ω_C_* values in HGTs were lower than the adaptive convergence events (Fig. 1E). By contrast, *C/D* and *dN_C_* showed values higher in HGTs than in the adaptive convergence events. Together with the simulations, these results show that the consideration of synonymous substitutions is essential for the accurate detection of molecular convergence in the presence of phylogenetic error and that *ω_C_* outperforms current alternative methods.

### Temporal variation of convergence rates

The probability of protein convergence decreases over time, with intramolecular epistasis among amino acid residues considered to be a primary biological source of such an evolutionary pattern (Goldstein et al., 2015; Zou and Zhang, 2015a; Goldstein and Pollock, 2017). Indeed, over a long timescale, the environment around any given focal site changes through substitutions at other amino acid sites, thus altering which amino acid state at the focal site is suitable to maintain structure and function (Goldstein and Pollock, 2017; Pollock et al., 2012) (Fig. S4L). However, gene tree discordance due to biological and technical causes, including tree inference error, incomplete lineage sorting, introgression, HGT, and intralocus recombination, can create a false convergence signal that similarly decreases with the time since branches separated (Mendes et al., 2016, 2019) (Fig. 1A and Fig. S1A). While the analysis of the mitochondrial genome (Goldstein et al., 2015) would not have been confounded by recombination-mediated mechanisms, other factors would have as great an influence as for nucleus-encoded genes. Nevertheless, all of the above problems would produce false convergence signals equally in synonymous and nonsynonymous substitutions via errors in the phylogenetic tree topology; therefore, *ω_C_* should be a natural candidate to unbiasedly evaluate whether convergence rates in nucleus-encoded genes also decrease with time.

We obtained 21 vertebrate genomes covering a range from fish to humans (Fig. 2A and Fig. S7A) and calculated *ω_C_* for all independent branch pairs in 16,724 orthogroups classified by OrthoFinder (Emms and Kelly, 2015, 2019). CSUBST completed the analysis even for the largest orthogroup (OG0000000), containing 682 genes encoding zinc finger proteins and 901,636 independent branch pairs (alignment length including gaps: 31,665 bp). We obtained a total of 20,150,538 branch pairs from all orthogroups and further analyzed 2,349,515 branch pairs with at least one synonymous and nonsynonymous convergence (i.e., 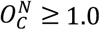 and 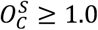). In all metrics (*C/D, dN_C_*, and *ω_C_*), protein convergence rates clearly decreased over time (approximated by inter-branch genetic distance) (Fig. 2B). Notably, we observed no such pattern for the rate of synonymous convergence (*dS_C_*), making it more likely that the diminishing protein convergence is caused by evolutionarily selected mechanisms (Goldstein et al., 2015; Zou and Zhang, 2015a). We also detected a similarly decreasing pattern in the rates of divergent substitutions over time, which does not contradict the effect of epistasis (Fig. S7B–C; Supplementary Text 7). Thus, the pattern of diminishing convergence remains a clear trend in recombining nucleus-encoded genes, even after correcting for the rate of synonymous convergence, and therefore is consistent with the action of intramolecular epistasis (Fig. S4L).

**Figure 2.**
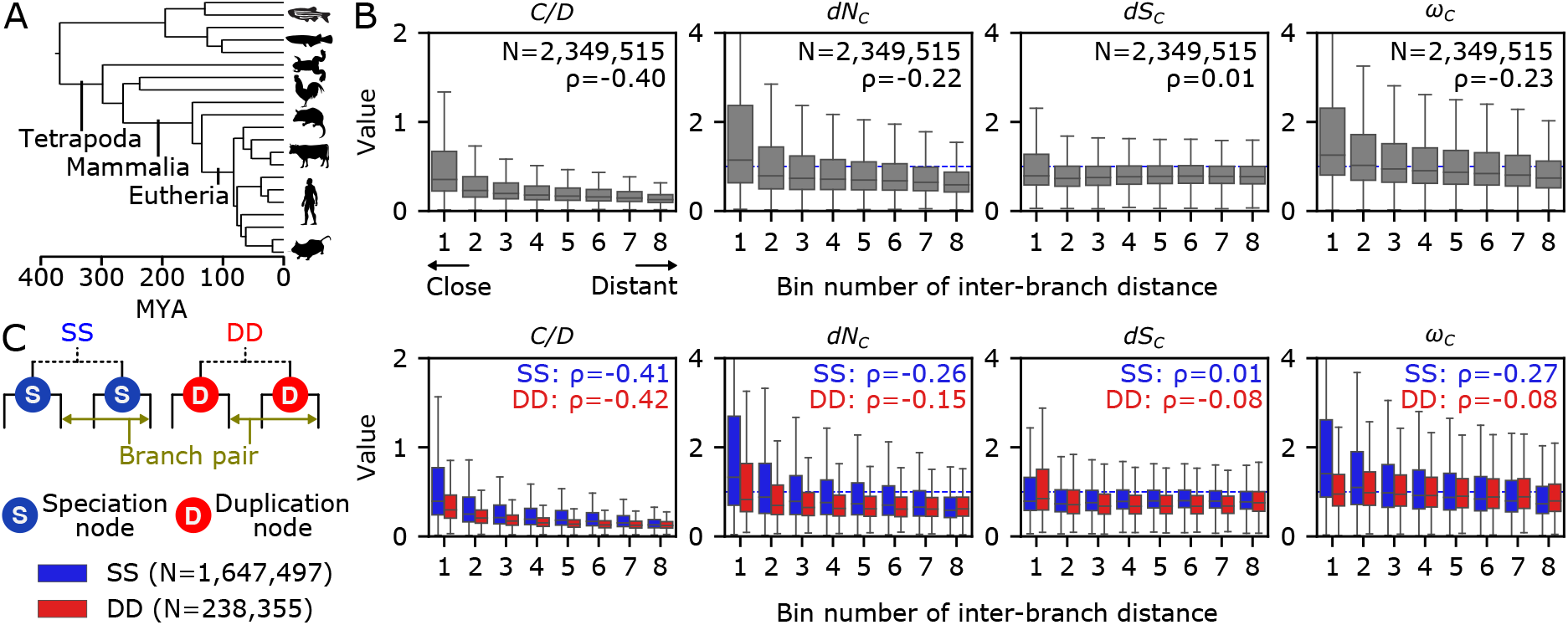
Biological variation of *ω_C_* in a genome-scale dataset. (**A**) Phylogenetic relationships of the selected species. See Fig. S7A for the complete phylogeny. The tree and divergence time estimates were obtained from timetree.org (Hedges et al., 2015). Some animal silhouettes were obtained from PhyloPic (http://phylopic.org). (**B**) Temporal variation of convergence rates. The numbers of branch pairs (N) and Spearman’s correlation coefficient (ρ) are shown. The bin range was determined to assign an equal number of branch pairs to each bin. To reduce the noise originating from branches where almost no substitutions occurred, branch pairs with both 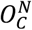 and 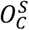 greater than 1 were analyzed (i.e., at least one convergent substitution each). (**C**) Convergence rates depending on gene duplications. Branch pairs were categorized into speciation events (SS) and branch pairs after two independent gene duplications (DD) according to the presence of preceding gene duplications in no or both branches, respectively. Branch pairs with one preceding duplication were excluded from the analysis. Dashed lines indicate the neutral expectation (=1.0).

### Gene duplication decreases convergence rates

Gene duplication generates new genetic building blocks (Conant and Wolfe, 2008) and elevates the rate of protein evolution (Fukushima and Pollock, 2020). However, it remains unknown whether substitution profile changes influence convergence rates following gene duplication. Convergent substitutions in duplicates may indicate convergent functional changes in independently duplicated genes, and our genome-scale dataset contains 90,028 duplication events, providing an excellent opportunity to address this question. If independent duplications in a family of genes tend to result in mutually similar derived pairs of proteins, the convergence rate should increase. Conversely, if the new proteins tend to move into a divergent sequence space in which they do not overlap, gene duplication would not increase convergence and may even decrease it. Accelerated non-adaptive change might not change the convergence rate if gene duplication only causes an increase in the rate of protein evolution without changing the substitution profiles. To distinguish these possibilities, we compared the convergence rates of branch pairs after two separate speciation (SS) events and branch pairs after two independent gene duplications (DD) (Fig. 2C). Strikingly, gene duplication significantly decreased convergence rates (*P* ≈ 0, *W* = 23.0, as determined by a two-sided Brunner–Munzel test; Fig. 2C). Again, the trend was evident in nonsynonymous convergence (*dN_C_*) but not in synonymous convergence (*dS_C_*), implying a relaxation in site-specific constraints or adaptive divergence in the duplicates. Notably, the effect of gene duplication was stronger in closely related branch pairs (i.e., smaller bin numbers in Fig. 2C), and the *ω_C_* distributions became progressively indistinguishable between SS and DD pairs with increasing inter-branch distance. The immediate drop of the convergence probability was consistent with the idea that gene duplication allows the new gene copies to explore a new sequence space, potentially involving natural selection. We note that this is an averaged trend across genes and does not exclude possible adaptive convergence in some genes. However, it is likely that such convergence, if it does exist, is masked by the opposing, predominant signal of relaxed or divergent constraints.

It is also noteworthy that the DD branch pairs show anomalously high synonymous convergence rates (*dS_C_*) in the smallest bin of genetic distance (bin 1 in Fig. 2C). This observation is probably due to the difficulty of locating gene duplication events in the phylogenetic tree, especially when sequences are not sufficiently diverged and lead to an extremely short branch length. Consistent with this idea, small genetic distances were associated with low branch supports in the DD branch pairs (Fig. S7D). Additionally, we detected similar anomalies in extremely distant branch pairs and attributed them to false orthogroup inference (Supplementary Text 8; Fig. S7E). These examples illustrate how various aspects of phylogenetic analysis can generate false patterns of convergence that are successfully captured by *dS_C_* and corrected for in *ω_C_*.

### Extracting a high-confidence set of convergent lineages

Discovering adaptive molecular convergence in genome-scale datasets, which may be translated into genotype-phenotype associations, has been challenging since it is a rare phenomenon and false positives are high (Foote et al., 2015; Thomas and Hahn, 2015; Zou and Zhang, 2015b). To examine whether the application of *ω_C_* can generate plausible hypotheses of adaptive molecular convergence, we analyzed the 21 vertebrate genomes (Fig. S7A). We first extracted the branch pairs with the top 1% of *C/D, dN_C_*, or *ω_C_* values with a cutoff for a minimum of three nonsynonymous and synonymous convergence (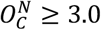 and 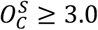) (Fig. 3A). The overlap between each set of branch pairs was moderate, with 1,348 branch pairs satisfying all three criteria out of 5,659 pairs with the top 1% *ω_C_* values.

**Figure 3.**
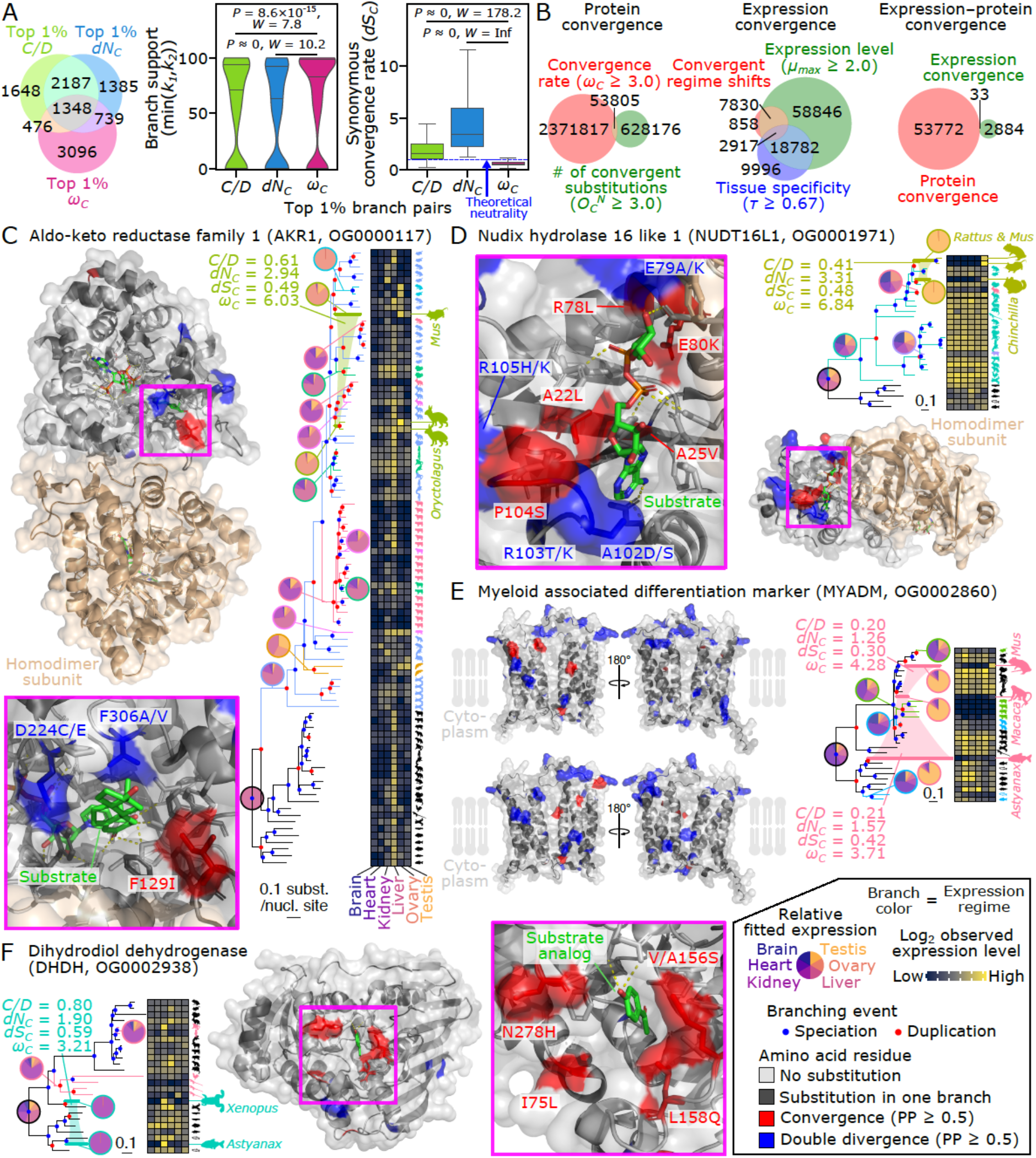
Joint convergence of gene expression patterns and protein sequences. (**A**) Comparison of convergent branch pairs obtained by different methods in the vertebrate dataset. Branch pairs with 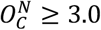 and 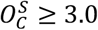 were analyzed. The Venn diagram on the left shows the extent of overlap between the top 1% convergent branch pairs. The violin plot in the middle shows the lower bootstrap support of the parental branches of the convergent branch pairs. The boxplot on the right compares the rate of synonymous convergence (*dS_C_*). The stochastic equality of data was tested by a two-sided Brunner–Munzel test (Brunner and Munzel, 2000). (**B**) Venn diagrams showing the extent of overlap between protein and expression convergence. Circles represent the sets of branch pairs. Shifts in tissue-specific expression regime were identified with the thresholds of expression levels (the maximum fitted SVA-log-TMM-FPKM among tissues (Fukushima and Pollock, 2020)) and tissue specificity (Yanai’s *τ* (Yanai et al., 2005)). (**C–F**) Examples of the likely adaptive joint convergence. Aldo-keto reductase family 1 (AKR1, **C**), Nudix hydrolase 16 like 1 (NUDT16L1, **D**), Myeloid associated differentiation marker (MYADM, **E**), and Dihydrodiol dehydrogenase (DHDH, **F**) are shown (see Fig. S9A for complete trees). Node colors in the trees indicate inferred branching events of speciation (blue) and gene duplication (red). The heatmap shows expression levels observed in extant species. The silhouettes signify the species (see Fig. S7A) that carries the gene, and the clades involved in the joint convergence are indicated with an enlarged size. The colors of branches and animal silhouettes indicate expression regimes. Among-organ expression patterns are shown as a pie chart for each regime. Branches involved in joint convergence are highlighted with thick lines, connected by the color of the expression regime, and annotated with convergence metrics. Localization of convergent and divergent substitutions on the protein structure is shown along with a close-up view of functionally important sites. The surface representation of each protein is overlaid with a cartoon representation. Convergent and divergent amino acid loci shown in Fig. S9 are highlighted in red and blue, respectively. Substrates and their analogs are shown as green sticks. Side chains forming the substrate-binding site are also shown as sticks. Note that these are the side chains in the protein from databases, so amino acid substitutions in the convergent lineages may result in distinct structures and arrangements. Site numbers correspond to those in the PDB entry or the AlphaFold structure (from **C** to **F**: 1Q13, 5W6X, AF-Q6DFR5-F1-model_v2, and 2O48). The silhouettes of *Astyanax mexicanus* and *Oreochromis niloticus* are licensed under CC BY-NC-SA 3.0 (https://creativecommons.org/licenses/by-nc-sa/3.0/) by Milton Tan (reproduced with permission), and those of *Anolis carolinensis* (by Sarah Werning), *Ornithorhynchus anatinus* (by Sarah Werning), and *Rattus norvegicus* (by Rebecca Groom; with modification) are licensed under CC BY 3.0 (https://creativecommons.org/licenses/by/3.0/).

To examine which metrics better enrich for likely adaptive convergence, we compared the topological confidence scores of the selected branches. If artifacts due to tree topology errors are included, low confidence branches should be enriched. Analysis of the bootstrap-based confidence values (Hoang et al., 2018; Minh et al., 2013) showed that *ω_C_* selects branch pairs with higher confidence than the other two metrics (Fig. 3A). Furthermore, we examined the synonymous convergence rate (*dS_C_*), which is not expected to be greater than the neutral expectation in the adaptive convergence, and established that only *ω_C_* satisfies such an assumption (Fig. 3A). These results indicate that *ω_C_* has excellent properties for finding adaptive protein convergence in genome-scale analyses.

### Identification of molecular convergence associated with a particular phenotype

As convergence metrics have been used to search for genes associated with phenotypes of interest, we next examined whether *ω_C_* might be used to discover candidate genes underlying phenotypic convergence. Here, we analyzed a pair of herbivorous animal lineages as an example: ruminants (cattle [*Bos taurus*] and red sheep [*Ovis aries*]) and rabbits (*Oryctolagus cuniculus*). Using minimum thresholds for the number of convergent amino acid substitutions 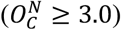 and protein convergence rate (**ω*_C_* ≥ 3.0), we obtained 352 candidate branch pairs in a genome-scale analysis of the 21 vertebrates (Table S5). By mapping the positions of substitutions onto known conformations of homologous proteins, we identified particularly compelling cases of likely adaptive convergence (Fig. S8). Examples included olfactory receptors in which convergent substitutions are located in the interior of the receptor barrel (ODORANT RECEPTOR 7A [OR7A], Olfactory Receptor Family 2 Subfamily M Member 2 [OR2M2], and OR1B1), where substitutions may change ligand preference associated with herbivorous behavior.

Similarly, the barrel-like structure of some solute carriers harbored convergent substitutions in their interior sides (Solute Carrier Family 5 Member 12 [SLC5A12], SLC51A, SLC22A, and SLC44A1), suggesting their involvement in the uptake or transport of plant-derived compounds. Among these, SLC51A (also known as Organic solute transporter α [OSTα]) may be a particularly attractive candidate. This protein plays a major role in bile acid absorption and, hence, in dietary lipid absorption (Ballatori et al., 2005). The convergence in SLC51A may be coupled with another convergent event detected in CYP7A1, a cytochrome P450 protein known to serve as a critical regulatory enzyme of bile acid biosynthesis (Chiang and Ferrell, 2020). CYP7A1 harbored two convergent substitutions in its substrate-binding sites (Fig. S8). While most herbivores secrete bile acids mainly in a glycine-conjugated form, ruminant bile is mostly in the form of taurine-conjugated bile acids, which remain soluble in highly acidic conditions (Noble, 1981). The predominance of taurine-conjugated forms is also observed in rabbits, depending on species and developmental stage (Hagey et al., 1998). Thus, convergence in these proteins may be related to such nutritional physiology.

Other examples of detected convergence included two convergent substitutions in the DNA-binding sites of a member of the zinc-finger protein family, which functions as a transcriptional regulator (Patel et al., 2018) (Fig. S8). Convergence in the substrate-binding sites of pancreatic elastase (Mulchande et al., 2007) and pancreatic DNase I (Weston et al., 1992) may be related to their specialized digestion (Fig. S8). In DNase I, amino acid sites exposed on the surface of protein structures displayed additional convergent substitutions that change the charge of their target amino acid residues (E124K, G172D, and H208N), possibly resulting in convergent changes in the biochemical properties of the protein, such as optimal pH, resistance to proteolysis, and posttranslational modifications. Consistent with this idea, bovine and rabbit DNase I proteins are known to be more resistant to degradation by pepsin than their homologs in other animals (Fujihara et al., 2012). Furthermore, E124K was shown to be important for the phosphorylation of bovine DNase I (Nishikawa et al., 1997). Other convergent substitutions will be promising candidates for future characterization. Taken together, these results show how our approach can detect genetic changes associated with phenotypes on the macroevolutionary scale.

### Exploratory analysis of as-yet-uncharacterized molecular convergence

We further exploited the 21 vertebrate genomes to examine whether *ω_C_* might be used to discover adaptive molecular convergence that may generate hypotheses of linked phenotypes. Since convergence at multiple levels of biological organization can provide strong evidence for adaptive evolution, we searched for simultaneous convergence in protein sequences and gene expression in an exploratory manner without a predefined hypothesis on convergently evolved genes and lineages. Using the same thresholds applied to the analysis of herbivores above (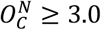 and *ω_C_* ≥ 3.0), we obtained 53,805 candidate branch pairs from all orthogroups (Fig. 3B).

Although this was an exploratory analysis in which all independent branch pairs were exhaustively analyzed, many studies of convergent evolution involve only a few groups of focal species. If such a research design is applied to this dataset (similar to the analysis of herbivores), the number of detected branch pairs will be much smaller. For example, because there are 538 independent branch pairs in the species tree, on average 100 cases of protein convergence will be obtained in our genome-scale dataset for any particular analysis of two groups of species.

To detect convergent gene expression evolution, we employed the amalgamated transcriptomes for six organs in the 21 vertebrate species (Fukushima and Pollock, 2020). Using this previously published dataset, we subjected curated gene expression levels (SVA-log-TMM-FPKM) to multi-optima phylogenetic Ornstein-Uhlenbeck (OU) modeling, in which expression evolution is inferred as regime shifts of estimated optimal expression levels (Khabbazian et al., 2016). Phylogenetic positions and the numbers of expression evolution were determined by a LASSO-based algorithm with Akaike Information Criterion, which was also used for finding convergent shifts toward similar optimal values. In total, we detected 12,017 cases of expression convergence in 4,308 orthogroups (Fig. 3B). Setting the thresholds for gene expression specificity at *τ* ≥ 0.67 (Yanai et al., 2005) and expression levels at *μ_max_* ≥ 2.0 (the maximum value of fitted SVA-log-TMM-FPKM) (Fukushima and Pollock, 2020), we obtained a set of 2,917 high-confidence branch pairs for potentially adaptive convergence of expression patterns.

By taking the intersection of protein convergence and expression convergence, we discovered 33 cases of potentially adaptive joint convergence of expression patterns and protein sequences in 31 orthogroups (Fig. 3B; Table S6). Gene duplication was frequently associated with joint convergence, with at least one branch experiencing gene duplication in 23 out of the 33 branch pairs (*P* = 3.11×10^−25^, *χ*^2^ = 107.7, *χ*^2^ test of independence). While gene duplication generally reduced the convergence rate, as discussed earlier (Fig. 2C), some of the independently generated duplicates may tend to evolve into the same sequence space when similar expression evolution takes place. Convergence of testis-specific genes was most frequently observed (19/33 orthogroups) and significantly enriched (*P* = 1.36×10^−31^, *χ*^2^ = 136.8, *χ*^2^ test of independence). The mechanism by which the testis serves as a major place for functional evolution of duplicated genes has been explained by several factors, including the ease with which expression is acquired in spermatogenic cells (Kaessmann, 2010; Kleene, 2005). This phenomenon is called the out-of-the-testis hypothesis, and our results suggest that predictable protein evolution may be enriched in this evolutionary pathway.

To infer the functional effect of convergent amino acid substitutions, we mapped the positions of substitutions onto known conformations of homologous proteins. Strikingly, we observed convergently evolved proteins where clusters of substitutions are localized to functionally important sites. They included members of aldo-keto reductase family 1 (AKR1), which play essential roles in steroid metabolism (Rižner and Penning, 2014). The OU analysis revealed that *AKR1* acquired preferential expression in the ovary after repeated lineage-specific duplications in rabbits and mice (*Mus musculus*) (Fig. 3C). Among the paired substitutions in the two lineages, F129I (convergence) and F306A/V (double divergence) located to the positions that delineate the steroid-binding cavity (Fig. 3C). At residue 306, the size of the amino acid was shown by targeted mutagenesis to be important for catalytic promiscuity in rabbits (Couture et al., 2004). Similarly, D224C/E (double divergence) occurred in a loop that contributes to substrate specificity (Couture et al., 2004). These results suggest that the phenotypic change related to substrate specificity might have occurred not only in rabbits but also in mice and underscore how F129I, together with the other two convergence cases (N11S and T/S289P, Fig. S9A), should be a major target for future characterization.

Similarly, *nudix hydrolase 16-like 1* (*NUDT16L1*, also known as *Tudor-interacting repair regulator* [*TIRR*]), which is involved in cell migration (Gunaratne et al., 2011) and whose encoded protein binds to RNA and P53-binding protein 1 (53BP1) (Botuyan et al., 2018), showed lineage-specific duplications in chinchillas (*Chinchilla lanigera*) and another rodent lineage connected to mice and rats (*Rattus norvegicus*) (Fig. 3D). The duplication events were followed by convergent regime shifts that resulted in testis-specific expression. The expression evolution was coupled with convergent substitutions in the protein sites corresponding to the substrate-binding pocket of the de-ADP-ribosylating homolog NUDT16 (Thirawatananond et al., 2019; Zhang et al., 2020). Protein convergence linked to testis-specific expression was also observed in *myeloid-associated differentiation marker* (*MYADM*), which encodes a transmembrane protein that localizes to membrane rafts (Aranda et al., 2011), regulates eosinophil apoptosis through binding to Surfactant protein A (SP-A) (Dy et al., 2021), and participates in cell proliferation and migration (Sun et al., 2016). This orthogroup showed joint convergence in two pairs of branches, in both of which the convergent amino acid substitutions were almost entirely confined to one side of the transmembrane domains (Fig. 3E), suggesting altered interactions with other molecules through this portion of the protein.

Finally, an orthogroup of dihydrodiol dehydrogenase (DHDH) showed joint convergence of expression and proteins (Fig. 3F). Possible physiological roles of this enzyme included the detoxification of cytotoxic dicarbonyl compounds, such as 3-deoxyglucosone derived from glycation (Nakayama et al., 1991; Sato et al., 1993). Although the domain structure of proteins was well conserved among species (Fig. S9A), the gene expression patterns of the encoding genes tended to vary. *DHDH* is known to show distinct tissue-specific expression patterns in mammals: kidney in monkeys (*Macaca mulatta*) (Nakagawa et al., 1989), kidney and liver in dogs (*Canis lupus*) (Sato et al., 1994), liver and lens in rabbits (Arimitsu et al., 1999), and various tissues in pigs (*Sus scrofa*) (Nakayama et al., 1991). Our amalgamated transcriptomes showed largely consistent species-specific expression patterns (Fig. 3F). The OU analysis recovered four lineage-specific regime shifts categorized into two pairs of convergent expression evolution. One of them, the convergence of gene expression that occurred between frogs (*Xenopus*) and the blind cave fish (*Astyanax*), which diverged approximately 435 million years ago (Hedges et al., 2015), is characterized by kidney-specific expression. The *Xenopus* gene ENSXETG00000033613 appeared to have arisen from a more widely expressed ancestral gene after a lineage-specific gene duplication. By contrast, the *Astyanax* gene ENSAMXG00000005808 may have acquired kidney-specific expression without any detectable duplication. In this branch pair, we detected a protein convergence rate that cannot be explained by neutral evolution, with a convergence of five amino acid sites (Fig. S9A). These convergent substitutions localized around the active site, while we did not observe such a trend for the double divergence (Fig. 3F). This result suggests that the convergent substitutions may have occurred adaptively to change ancestral catalytic function.

DHDH has a broad substrate specificity for carbonyl compounds. This protein oxidizes *trans*-cyclohexanediol, *trans*-dihydrodiols of aromatic hydrocarbons, and monosaccharides including D-xylose, while it reduces dicarbonyl compounds, aldehydes, and ketones (Sato et al., 1994). Its active site is predominantly formed by hydrophobic residues, suggesting their role in catabolizing aromatic hydrocarbons (Carbone et al., 2008b, 2008a). Notably, the convergent substitutions in the substrate-binding sites tended to increase amino acid hydrophobicity (Fig. S9B), suggesting that the remodeling of the active site may have led to the acquisition of new substrates in *Xenopus* and *Astyanax*.

In summary, *ω_C_* was not only robust against phylogenetic errors, outperforming other methods in simulation and empirical data, but also allowed us to discover plausible adaptive convergence from a genome-scale dataset without a pre-existing hypothesis. Molecular convergence revealed by our exploratory analysis will provide a basis for understanding overlooked phenotypes that protein evolution led to in corresponding lineages.

### Heuristic detection of highly repetitive adaptive convergence

Convergent events observed on even more than two independent lineages are exceptionally good signals of adaptive evolution, if they exist, because three or more combined convergences should be extremely rare in random noise. Conventionally, convergence in more than two branches has been analyzed as multiple pairwise comparisons for which there is a prior hypothesis of convergence. The difficulty in analyzing higher-order combinatorial substitutions without specific prior hypotheses lies in the need to explore a vast combinatorial space that exponentially expands as the number of branches to be combined (*K*) increases. For example, an evenly branching tree with 64 tips has 7,359 independent branch pairs (i.e., at *K* = 2), but the number of branch combinations exponentially increases to 333,375 and 6,976,859 in triple (*K* = 3) and quadruple (*K* = 4) combinations, respectively, making it impractical to exhaustively search highly repetitive convergence even in a single phylogenetic tree when a hypothesis on focal lineages is unavailable.

To overcome this limitation, we developed an efficient branch-and-bound algorithm (Land and Doig, 1960) that progressively searches for higher-order branch combinations (Fig. 4A and Fig. S10A). For the performance evaluation, we used the PEPC tree (Fig. 4B) because it has repeated adaptive convergence for its use in C_4_ photosynthesis (Fig. 1E). While the exhaustive search required 156 minutes with *K* = 3 to analyze 307,432 branch combinations using two central processing units (CPUs), our branch-and-bound algorithm required only 21 seconds. At *K* = 4, the exhaustive search completed within a practical time by using 16 CPUs (46 hours for nearly 8 million combinations) but failed to complete at *K* = 5 (152 million combinations). By sharp contrast, the heuristic search took about 5 minutes for the entire analysis, of which the higher-order analysis with *K* ranging from 3 to 6 took only about 1 minute to analyze as few as 390 combinations with two CPUs (Table S7).

**Figure 4.**
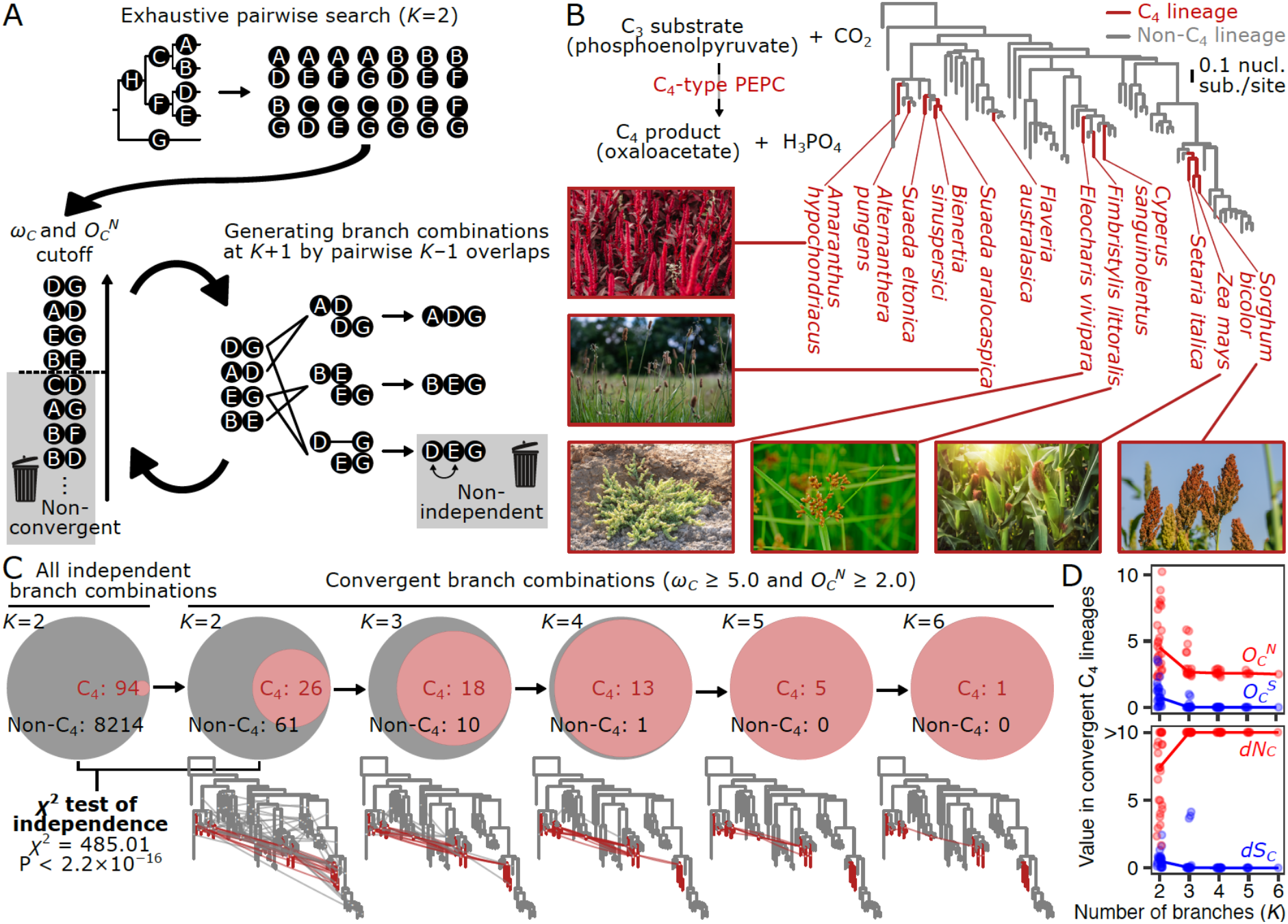
Heuristic search of higher-order branch combinations for adaptive protein convergence. (**A**) Branch-and-bound algorithm for higher-order branch combinations. This method explores the higher-order combinatorial space until there are no more convergent branch combinations. (**B**) The maximum-likelihood phylogenetic tree of phosphoenolpyruvate carboxylases (PEPCs) in flowering plants. The catalytic function of PEPC, which is crucial in C_4_ photosynthesis, is illustrated. Photographs of representative C_4_ photosynthetic lineages are shown. The photograph of *Suaeda aralocaspica* is reproduced from the literature (Wang et al., 2019). The bar indicates 0.1 nucleotide substitutions per nucleotide site. The complete tree is shown in Fig. S5. (**C**) Higher-order convergence enriches C_4_-type PEPCs. The Venn diagrams show the proportion of convergent branch combinations of C_4_-type and non-C_4_-type lineages (red and gray, respectively). Branch combinations containing both were included in non-C_4_. In the phylogenetic trees, convergent branch combinations are shown as edges connecting branches. (**D**) Improvement of the signal-to-noise ratio in higher-order branch combinations. The line graph shows the median values of the total probabilities (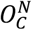 and 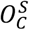) and the rates (*dN_C_* and *dS_C_*) of nonsynonymous and synonymous convergence in the convergent branch combinations of C_4_ lineages. Points correspond to branch combinations.

The analyzed tree covered nine independent origins of C_4_-type PEPC, and the corresponding branch pairs of C_4_ lineages accounted for 1.1% of all possible pairs (94/8,308). Convergent branch pairs defined by a threshold (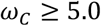 and 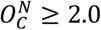) enriched for the C_4_ lineages at *K* = 2 (29.9%, 26/87; Fig. 4C). The convergence of non-C_4_ lineages (61/87, including pairs of C_4_ and non-C_4_ branches) can be interpreted as false positives or adaptive convergence associated with other currently unknown functions. The subsequent higher-order analysis resulted in the discovery of highly repetitive convergence in combinations of as many as six branches (i.e., *K* = 3 to *K* = 6). As the order increased, the lineages of C_4_-type PEPCs rapidly predominated and accounted for all the combinations detected at *K* ≥ 5 (Fig. 4C), even though the heuristic algorithm was not given any information about the C_4_ lineages.

In the higher-order C_4_ branch combinations, the detected convergence events were almost entirely nonsynonymous 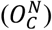, while synonymous convergence 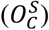 was negligible (Fig. 4D). As a result, the rate of synonymous convergence (*dS_C_*) quickly approached zero (Fig. 4D). Notably, the higher-order convergent substitutions were located at functionally important protein sites. In the convergent branch combinations with *K* = 6, we identified three amino acid sites with a joint posterior probability of nonsynonymous convergence greater than 0.5: V627I, H665N, and A780S (Fig. S10B–D). The H665N substitution generates a putative N-glycosylation site that may be important for protein folding (Christin et al., 2007). The A780S substitution, for which the signature of positive selection had been detected previously (Besnard et al., 2009; Hermans and Westhoff, 1992; Poetsch et al., 1991), has been shown to change the enzyme kinetics related to the first committed step of C_4_ carbon fixation (Bläsing et al., 2000; DiMario and Cousins, 2019; Engelmann et al., 2002) and is therefore considered a diagnostic substitution of C_4_-type PEPC (Besnard et al., 2009; Christin et al., 2007). The third substitution, C627I, might be a good focus for future experimentation. These results demonstrate that higher-order analysis can substantially increase the signal-to-noise ratio in convergence analysis when there is repeated selective pressure to evolve similar biochemical functions.

## Discussion

In this study, we introduced a measure of convergent protein evolution, *ω_C_*, designed to account for false signals due to phylogenetic error. We showed, through simulation and analysis of real biological data, that *ω_C_* mostly eliminates false positives without reduction in power to detect true signals. We also developed an approach to estimate the rates of highly repetitive convergence (i.e., on more than two lineages) fully accounting for phylogenetic combinatorics and demonstrated that the specificity of *ω_C_* increases further in the higher-order analysis. Because of its improved accuracy, *ω_C_* should further drive macroevolutionary analyses where uncorrected measures have been used to identify responsible genotypes for particular phenotypes in a way similar to genome-wide association studies (GWASs). As in GWAS-identified alleles (or genes in gene-level association tests (Wang et al., 2021)), genes with excess convergence serve as clues to study macroevolutionary traits for which the molecular basis is unknown (Fig. 5). Furthermore, the accuracy of *ω_C_* even allows exploratory analysis (Fig. 5), as demonstrated here in vertebrate genomes (Fig. 3). By conducting a genome-wide search of convergent branch combinations, we detected signatures of likely adaptive convergence, which leads to hypothesis generation on responsible phenotypes. This outcome was possible because *ω_C_*, unlike *P*-values from GWASs, does not require phenotypic traits as input. Convergently evolved genes identified by exploratory analysis will, in turn, lead to the discovery of overlooked phenotypes through future experimentation.

**Figure 5.**
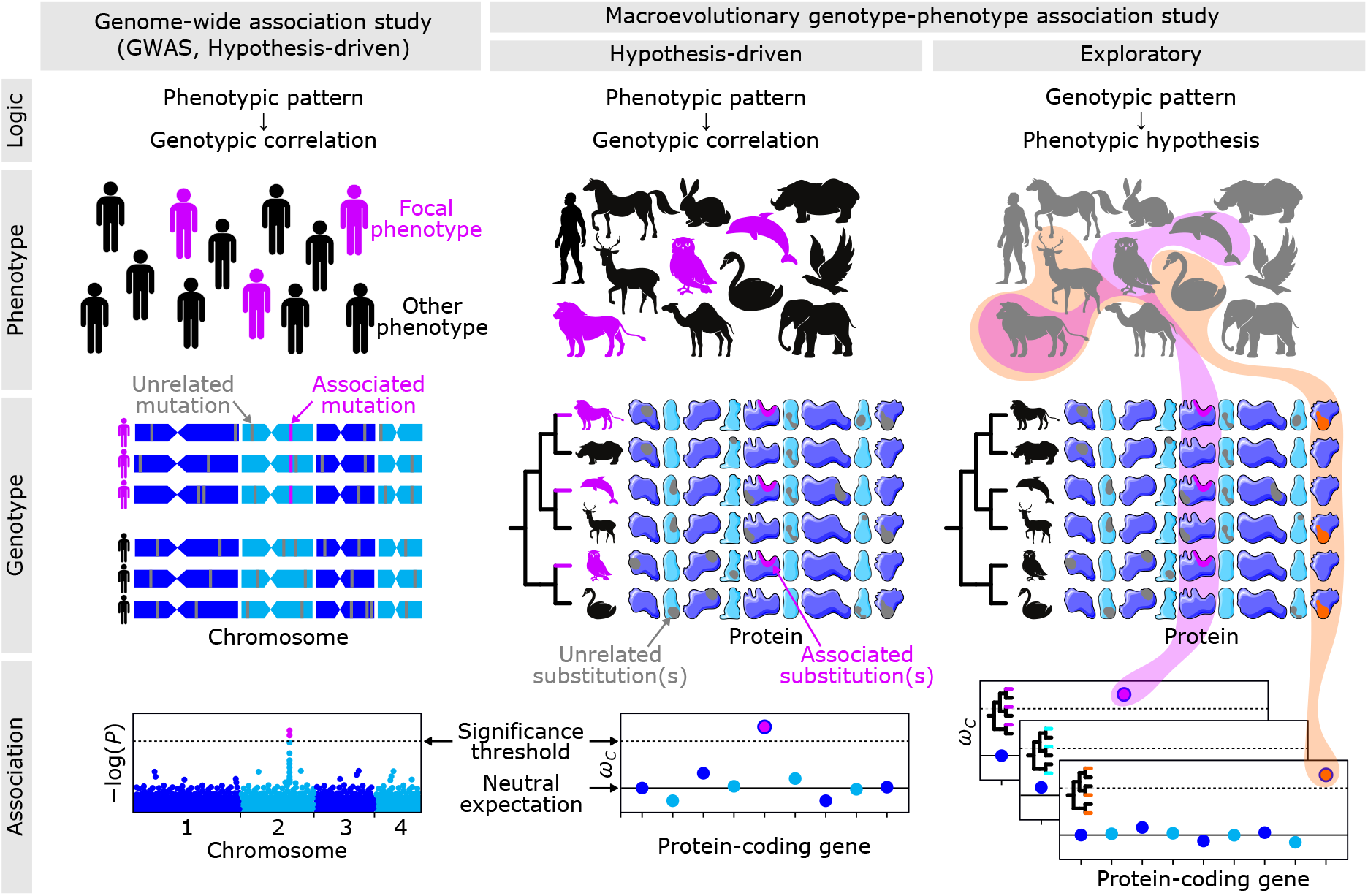
Analysis of the genotype-phenotype association within and between species. The proposed method improves the accuracy of the hypothesis-driven approach in the macroevolutionary scale and enables exploratory approaches. Note that for visualization purposes, the number of individuals and species shown here is smaller than the actual number required for analysis. The icons of proteins are licensed under CC BY 3.0 (https://creativecommons.org/licenses/by/3.0/) by Smart Servier Medical Art.

Although *ω_C_* is a powerful means to detect convergence while removing the effect of phylogenetic error, there are other sources of stochastic error that can mask small signals. We successfully captured multiple known convergence events here, even with only two or three amino acid substitutions involved in small proteins (Fig. 1E and Table S3). However, a convergent amino acid substitution at a single site in only two lineages may not reliably be identified as resulting from adaptation rather than random homoplasy, by *ω_C_* or any other measure. Therefore, the number of observed nonsynonymous convergence 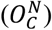 should always be considered in addition to the phylogenetic error-corrected convergence rate (**ω*_C_*), especially in a genome-scale screening with only two or three focal lineages. If many amino acid sites and/or many separate lineages are involved, true convergence is, in general, more easily detected.

Protein convergence has attracted a great deal of attention for its potential to associate long-term genotypic variation with phenotypic change, from its first discovery (Stewart et al., 1987), subsequent theoretical development (Castoe et al., 2009; Zhang and Kumar, 1997), the first claim of genome-wide detection (Parker et al., 2013), to recent findings that highlighted epistatic effects (Goldstein and Pollock, 2017; Goldstein et al., 2015; Zou and Zhang, 2015a, 2017) and technical difficulties (Foote et al., 2015; Mendes et al., 2016, 2019; Thomas and Hahn, 2015; Zou and Zhang, 2015b). Other types of convergence at the molecular level beyond amino acid substitutions have also been considered, including convergent shifts of site-wise substitution profiles (Rey et al., 2018), convergent shifts of evolutionary rates (i.e., number of substitutions per time regardless of the amino acid state or substitution profile) (Kowalczyk et al., 2019), convergent rate shifts of noncoding elements (Hu et al., 2019), convergent gene losses (Hiller et al., 2012; Prudent et al., 2016), convergent losses of noncoding elements (Marcovitz et al., 2016), and functional enrichments of convergently evolved loci (Marcovitz et al., 2019). Using transcriptome amalgamation, which integrates multi-species gene expression data in a comparable manner (Fukushima and Pollock, 2020), we developed a means to detect convergence in gene expression levels and to correlate the obtained results with protein convergence rates. Further integration of these methods will allow us to examine how well convergent patterns correlate across multiple hierarchies of biological organizations. Such analysis will provide a quantitative perspective of the extent to which evolution at one hierarchical level causes predictable changes in another.

Although it is well established that phenotypes are associated with genotypes, the genetic basis for particular convergently evolved phenotypes may arise from distinct, non-convergent genetic changes (Concha et al., 2019; Natarajan et al., 2016). These specific cases may sometimes occur because of convergent mechanisms, such as the use of similar but not identical amino acids, and the use of similar changes at adjacent residues in the protein structure (Castoe et al., 2008). The accumulation of knowledge about which mutations are repeatedly selected and which are not during convergent evolution may provide insight into the evolvability and constraints that govern the diversification of organisms.

While some evolutionary innovations may be unique, many traits arose convergently (Vermeij, 2006). Fascinating examples not mentioned above include endothermy, hibernation, burrowing, diving, venom injection, electrogenic organs, eusociality, anhydrobiosis, bioluminescence, biomineralization, plant parasitism, mycoheterotrophy, and multicellularity. In the past, the observation of similar phenotypes in multiple species led to the theory of evolution by natural selection (Darwin, 1859). The analysis of protein sequences in multiple species gave rise to the formulation of the nearly neutral theory of molecular evolution (Kimura, 1968; Ohta, 1973). Likewise, cross-species genotype-phenotype associations illuminated through the analysis of molecular convergence, coupled with experimental evaluation of mutational effects (Supplementary Text 9), may lead to new conceptual frameworks on the constraint and adaptive changes at the molecular level that drive phenotypic change among species.

## Methods

### Simulated codon sequence evolution

With the input phylogenetic tree (Fig. 1C), codon sequences of specified length (500 codons) were generated with the ‘simulate’ function of CSUBST (https://github.com/kfuku52/csubst), which internally utilizes the python package pyvolve for simulated sequence evolution (Spielman and Wilke, 2015). An empirical codon substitution model with multiple nucleotide substitutions (Kosiol et al., 2007) was adjusted with observed codon frequencies (ECMK07+F) in the vertebrate genes encoding phosphoglycerol kinases (PGKs, available from the ‘dataset’ function of CSUBST). The conventional *ω* (*dN/dS*) was set to 0.2. In the Convergent scenario, 5% of codon sites were evolved convergently in focal lineages (the pair of terminal branches in Fig. 1C). At convergent codon sites, the frequency of nonsynonymous substitutions to codons encoding a single randomly selected amino acid was increased so that nonsynonymous substitutions to the selected codons accounted for approximately 90% of the total. This operation increases the probability of amino acid convergence without changing relative frequencies among synonymous codons. The site-specific substitution rate at convergent codon sites was also doubled (i.e., *r* = 2), and a higher nonsynonymous/synonymous substitution rate ratio was applied (i.e., *ω* = 5) to mimic adaptive evolution. The simulation parameters for the other scenarios are summarized in Table S2. For the Random scenario, randomized trees were generated in 1,000 simulations with the ‘shuffle’ function of NWKIT v0.10.0 and the --label option (https://github.com/kfuku52/nwkit).

### Animal gene sets

A dataset of amalgamated cross-species transcriptomes (Fukushima and Pollock, 2020) was generated for 21 vertebrate genomes in Ensembl 91 (Yates et al., 2016) (Table S8). To ensure compatibility, the same versions of protein-coding sequences were also used for the convergence analysis. Completeness of genome assembly was evaluated using BUSCO v4.0.5 (Simão et al., 2015) with the single-copy gene set of ‘tetrapoda_odb10’ (Table S8). A species phylogenetic tree previously downloaded from TimeTree (Hedges et al., 2006) was used (Fukushima and Pollock, 2020). Orthogroups were classified by OrthoFinder v2.4.1 (Emms and Kelly, 2015, 2019). Orthogroups containing more than three genes were analyzed further. During the analysis of this dataset, a protein size–dependent change in measured convergence rates was observed (Fig. S11) but was determined to be an artifact; *ω_C_* was shown to be more robust to the bias than the other metrics (Supplementary Text 10).

### Sequence retrieval from public databases

Gene sets for previously confirmed cases of molecular convergence and horizontal gene transfer events (HGTs) were generated based on previous reports with increased taxon sampling (Table S3; Supplementary Text 11). With GenBank accession numbers for ATPalpha1, Prestin, PEPC, and PCK homologs (Supplementary Dataset), coding sequences (CDSs) were retrieved using the ‘accession2fasta’ function of CDSKIT. Lysozyme sequences were downloaded as GenBank files from NCBI and were converted to fasta files with the ‘parsegb’ function of CDSKIT. For the retrieval of the mitochondrial genome, a custom python script was used to select balanced numbers and lineages of foreground and background species (Supplementary Dataset). Orthogroup CDS files for og3737 (leucine-tRNA ligase), og9103 (pentatricopeptide repeat protein), and og9298 (pentatricopeptide repeat protein) for the HGT events in *Cuscuta* were obtained from a previous report (Yang et al., 2019b), and genes leading to unrealistically long branches were excluded. HGTs in the other parasitic lineage Orobanchaceae were also analyzed in the same report, but HGTs in *Cuscuta* were used for performance evaluation because the donor lineage was unequivocal in several genes.

### Sequence retrieval from plant gene sets

Gene sets were downloaded from public databases for the retrieval of CDSs encoding digestive enzyme homologs (Table S8). Transcriptome assemblies were used as a part of gene sets. For *Drosera adelae, Nepenthes* cf. *alata*, and *Sarracenia purpurea*, previously assembled transcriptomes were used (Fukushima et al., 2017). The transcriptome assembly of *Rhododendron delavayi* was generated from publicly available RNA-seq data (NCBI BioProject ID: PRJNA476831) with Trinity v2.8.5 (Grabherr et al., 2011) after pre-processing with fastp v0.20.1 (Chen et al., 2018) (Supplementary Dataset). Subsequently, open reading frames (ORFs) were obtained with TransDecoder v5.5.0 (https://github.com/TransDecoder/TransDecoder). The longest ORFs among isoforms were extracted with the ‘aggregate’ function of CDSKIT v0.9.1 (https://github.com/kfuku52/cdskit). The completeness of assembly was evaluated using BUSCO scores with the single-copy gene set of ‘embryophyta_odb10’ (Table S8). Finally, digestive enzyme homologs were retrieved by TBLASTX v2.9.0 searches against all gene sets with an E-value cutoff of 0.01 and >50% query coverage (Camacho et al., 2009).

### Characterization of protein-coding sequences

Coding sequences were used for RPS-BLAST v2.9.0 searches (Camacho et al., 2009) against Pfam-A families (El-Gebali et al., 2019) (released on April 30, 2020) with an E-value cutoff of 0.01 to obtain protein domain architectures. The numbers of transmembrane domains were predicted by TMHMM v2.0 (Krogh et al., 2001). The numbers of introns in protein-coding sequences were extracted from GFF files downloaded from Ensembl. Further gene annotations were obtained using Trinotate v3.2.1 (https://github.com/Trinotate/Trinotate.github.io/wiki).

### Plant species tree

Orthogroup classification was performed with OrthoFinder v2.4.1 (Emms and Kelly, 2019). Stop codons and ambiguous codons were masked as gaps using CDSKIT. In-frame multiple sequence alignments of single-copy orthologs were generated by MAFFT v7.455 with the --auto option (Katoh and Standley, 2013) and tranalign in EMBOSS v6.6.0 (Rice et al., 2000). Ambiguous codon sites were then removed by ClipKIT v0.1.2 with the default parameters (Steenwyk et al., 2020). After the concatenation of trimmed sequences, a maximum-likelihood phylogenetic tree was reconstructed by IQ-TREE v2.0.3 with the GTR+G nucleotide substitution model (Minh et al., 2020; Nguyen et al., 2015). The tree was rooted using *Amborella trichocarpa* as an outgroup. The divergence time of the species tree was estimated using mcmctree in the PAML package v4.9 (Yang, 2007). The priors and parameters were chosen according to the mcmctree tutorial (http://abacus.gene.ucl.ac.uk/software/paml.html). Fossil calibrations were adopted from a previous study (Zhang et al., 2017).

### In-frame codon sequence alignment

Retrieved coding sequences were formatted into in-frame sequences using the ‘pad’ function of CDSKIT. Stop codons and ambiguous codons were replaced with gaps with the ‘mask’ function of CDSKIT. Amino acid sequences from translated coding sequences were aligned using MAFFT with the --auto option (Katoh and Standley, 2013), trimmed with ClipKIT with default parameters, and reverse-translated with the ‘backtrim’ function of CDSKIT. Gappy codon sites were excluded with the ‘hammer’ function of CDSKIT.

### Phylogenetic tree reconstruction

The gene tree was first reconstructed using IQ-TREE with the general time-reversible (GTR) nucleotide substitution model and four gamma categories of among-site rate heterogeneity (ASRV). To suppress branch attraction in the trees containing HGTs, topological constraints consistent with species classification were generated from the NCBI Taxonomy (Schoch et al., 2020) using the ‘constrain’ function of NWKIT and used for tree search. Ultrafast bootstrapping with 1,000 replicates was performed to evaluate the credibility of tree topology (Minh et al., 2013) with further optimization of each bootstrapping tree (-bnni option) (Hoang et al., 2018). To improve tree topology, some datasets were subjected to phylogeny reconciliation with the species tree using GeneRax v1.2.2 (Morel et al., 2020) (Table S3). Branching events in gene trees were categorized into speciation or gene duplication by a species-overlap method (Huerta-Cepas et al., 2007). *Arabidopsis thaliana* orthologs in each clade were inferred from the tree topology. Minor differences in the methods applied to each dataset, from sequence retrieval to phylogenetic analysis, are summarized in Table S3.

### Detecting convergent expression evolution

Using the dated species tree and rooted gene trees as inputs, the divergence time of individual gene trees was estimated by RADTE (https://github.com/kfuku52/RADTE) as described previously (Fukushima and Pollock, 2020). Evolution of gene expression levels (SVA-log-TMM-FPKM) (Fukushima and Pollock, 2020) in brain, heart, kidney, liver, ovary, and testis samples was modeled on the dated gene tree with phylogenetic multi-optima Ornstein-Uhlenbeck models (i.e., Hansen models (Hansen, 1997)) with the ‘estimate_shift_configuration’ function in the R package *l*1ou v1.40 (Khabbazian et al., 2016) as described previously (Fukushima and Pollock, 2020). Convergent regime shifts were then detected as multiple regime shifts that lead to similar expression levels, as judged by the ‘estimate_convergent_regimes’ function (Khabbazian et al., 2016).

### Classification of combinatorial substitutions

Combinatorial substitutions were collectively defined as substitutions at the same protein site that occur in multiple independent branches in a phylogenetic tree. When this occurs only in two branches, it is called a paired substitution. In unambiguous notation, we consider paired substitutions along two branches with the same specific state (spe), different states (dif), or any state (any) at the ancestral and derived nodes. The five combinatorial states that we discuss and that are frequently considered in the literature are paired substitutions (any→any), double divergence (any→dif), convergence (any→spe), discordant convergence (dif→spe), and congruent convergence (spe→spe) (Fig. S1C). Convergence is discussed throughout this report because it is of particular importance in testing evolutionary genotype-phenotype associations.

### Ancestral state reconstruction and parameter estimation

Our method estimates convergent substitution via ancestral reconstruction. Whereas ancestral amino acid reconstruction has been used in previous reports (Foote et al., 2015; Goldstein et al., 2015; Thomas and Hahn, 2015; Zou and Zhang, 2015a), here we used codon sequence reconstruction. Using the input phylogenetic tree and observed codon sequences, CSUBST internally uses IQ-TREE to estimate the posterior probabilities of ancestral sequences by the empirical Bayesian method (Minh et al., 2020). At the same time, the parameters used in CSUBST are estimated: codon equilibrium frequencies (*π_i_*), ASRV (*r_l_*), nonsynonymous per synonymous substitution ratio (*ω*), and transition per transversion substitution ratio (*κ*).

### Multidimensional array structures for substitution history

CSUBST stores the coding sequences and the reconstructed probable ancestral states in a three-dimensional array whose size is *M* × *L* × 61 for a phylogenetic tree with *M* nodes (excluding the root node) generated from an alignment of coding sequences with *L* codon sites, each of which can take a distribution of 61 different codon states (in the universal genetic code), excluding stop codons. We denote by *P_mlj_*(*X*|*D, θ*) the posterior probability of codon *X* for codon state *j* at site *l* on node *m*. The three-dimensional array for codon states is then converted to a four-dimensional array that stores the probability of substitutions with the size of *B* × *L* × 61 × 61, where *B* denotes the number of branches excluding the root branch. This array stores the posterior probability of substitution *P_blij_*(*S* |*D, θ*) for single substitution *S* from ancestral codon state *i* to derived codon state *j* for a codon site *l* in branch *b*. For a site *l* in branch *b* connecting ancestral node *n* with codon state *i* and descendant node *m* with codon state *j*, the posterior probability substitution matrix *P_ij_*(*S*|*D, θ*) is derived as

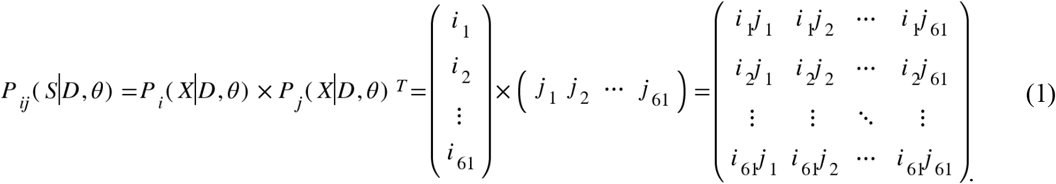

As the transition between the same codon state is not considered a substitution, the diagonal elements (*ij_i=j_*) are filled with 0. Although Equation 1 is an approximation that does not take into account the non-independence between nodes of a phylogenetic tree, we confirmed that the effect was negligible (Supplementary Text 12; Fig. S12). For efficient processing of nonsynonymous and synonymous substitution probabilities with the array operation of NumPy (Harris et al., 2020), the four-dimensional array is converted into a pair of five-dimensional arrays (*A^N^* and *A^S^* for nonsynonymous and synonymous substitutions, respectively) whose individual size is *B* × *L* × *G* × *I* × *J*, where codon states are grouped into *G* categories (Fig. S2A). Stored values range between 0 and 1, denoted by *P_blgij_*(*S*|*D*, *θ*), the probability of single substitution *S* from ancestral codon *i* to derived codon *j* (*i* ≠ *j*) in codon group *g* at site *l* of branch *b*, given the observed sequence data *D* and model parameters *θ* that include the phylogenetic tree. The elements in the array *A^N^* indicate *P_blgij_*(*S^N^*|*D*, *θ*), the probabilities of nonsynonymous substitutions (*S^N^*), whereas those in the array *A^S^* correspond to *P_blgij_*(*S^S^*|*D, θ*), the probabilities of synonymous substitutions (*S_S_*). In *A^N^*, a single 20×20 matrix records all the substitution probabilities, and therefore *G* = 1 and *I* = *J* = 20. Synonymous substitutions occur only between codons that code for the same amino acid. Since there are 20 different amino acids, *G* equals 20 in *A^S^*. In the case of the universal genetic code, the maximum number of codons encoding the same amino acid is six, for leucine, serine, and arginine, so *I* = *J* = 6. In the matrix corresponding to these three amino acids, all values are between 0 and 1, but for amino acids with a smaller number of codons, the out-of-range indices are filled with zero. Missing sites in the sequence alignment are also treated as zero. For simplicity, we explain the case where there is no missing site in the observed sequences and ancestral states in the following sections, but the implementation in CSUBST appropriately takes into account the missing sites by subtracting its numbers from *L* at every necessary step in individual branches or branch combinations.

### Tree rescaling

During the ancestral state reconstruction, IQ-TREE estimates the branch length as the number of nucleotide substitutions per codon site. Since our model requires the number of codon substitutions rather than the number of nucleotide substitutions, and since branch lengths are required separately for both synonymous and nonsynonymous substitutions, we obtained rescaled branch length *t_b_* of branch *b* as follows:

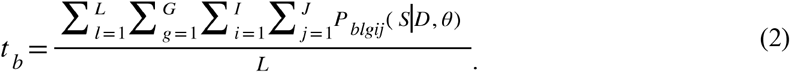

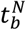 and 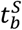 for nonsynonymous and synonymous substitutions were obtained with *P_blgij_*(*S^N^*|*D,θ*) and *P_blgij_*(*S^S^*|*D, θ*), respectively. For example, with the ECMK07+F+R4 model, the total branch lengths of the 21 vertebrate PGK tree before and after rescaling are 7.57 nucleotide-substitutions/codon-site and 7.20 codon-substitutions/codon-site (1.59 nonsynonymous and 5.62 synonymous codon substitutions per codon site).

### Observed number of combinatorial substitutions

The only true observations are the gene sequences of the extant species, and the posterior probabilities of ancestral sequences and codon substitutions are estimates. However, we refer to the posterior probabilities as “observations” (Zou and Zhang, 2015a) to unambiguously distinguish them from the expected values described in the next section. Here, we denote by *P_l_*(*S_C_*|*D, θ*) the probability of combinatorial substitution *S_C_* at codon site *l* given observed sequences *D* and model *θ*. The probabilities of nonsynonymous and synonymous combinatorial substitutions at site *l* are separately obtained as 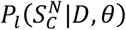 and 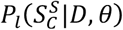, respectively, with the following equations:

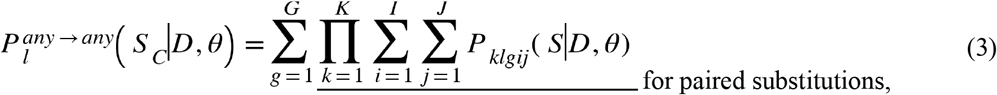

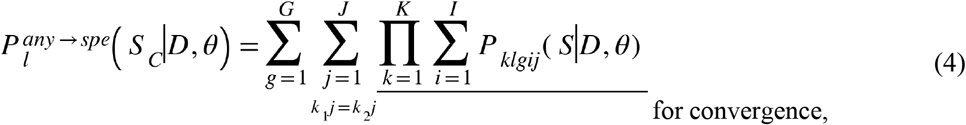

and

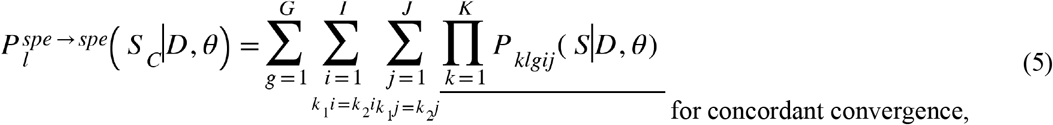

where *k* represents a branch of interest. We denote by *K* the degree of combinatorial substitutions or the number of branches to be compared. Because two branches are often compared in conventional convergence analysis, we explain here the case of *K* = 2. Array operations in the underlined parts of Equation 3 to Equation 5 are illustrated in Fig. S2B. The total probabilities of observed substitution pairs across sites in the branch pair are calculated as

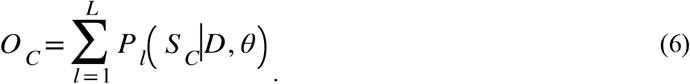

*O_C_* is separately obtained for nonsynonymous and synonymous combinatorial substitutions (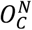 and 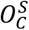, respectively). By definition (Fig. S1C), the values of *O_C_* for double divergence and discordant convergence are derived as follows at *K* = 2:

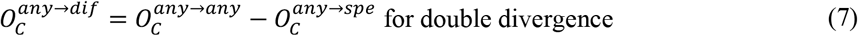

and

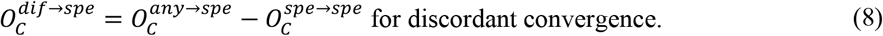

*C/D* (Goldstein et al., 2015) corresponds to 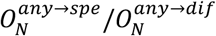 in our notation.

### Applying codon substitution models for the expectation of combinatorial substitutions

To estimate the rate of combinatorial substitutions, the observed number *O_C_* is contrasted with the expected number *E_C_*. *E_C_* is derived from codon substitution models in a way similar to the previous application of amino acid substitution models (Zou and Zhang, 2015a). The tested codon substitution models include the empirical models ECMK07 and ECMrest (Kosiol et al., 2007) and the mechanistic models MG (Muse and Gaut, 1994) and GY (Goldman and Yang, 1994). The same model was consistently used in the ancestral state reconstruction and in deriving the model-based expectations of combinatorial substitutions. In the method described below, empirical equilibrium codon frequencies, the rescaled branch length, and ASRV are also taken into account. In the empirical models, the codon substitution rate matrix *Q* is derived according to previous literature (Kosiol et al., 2007; Whelan and Goldman, 2001) as follows:

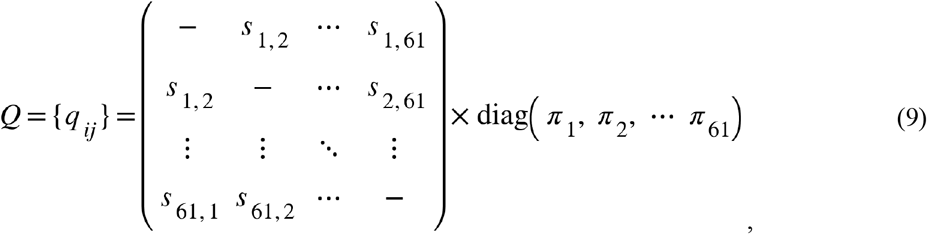

where *s_i,j_* denotes the exchangeabilities of codon pairs *i* and *j* (*s_ij_* = *s_ij_*), and *π_i_* represents the equilibrium frequencies of 61 codons estimated from the input alignment. In the mechanistic models, mechanistic substitution parameters are used instead of the exchangeabilities. In the MG model, *q_ij_* is obtained with *π_i_* and nonsynonymous per synonymous substitution ratio *ω*, whereas transition per transversion substitution ratio *κ* is also taken into account in the GY model. *Q* is then rescaled as

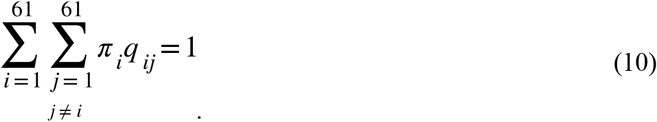

Finally, the diagonal elements of *Q* are completed as

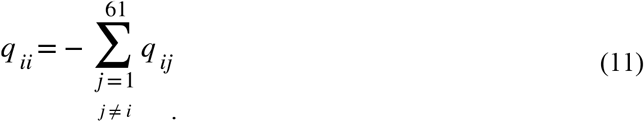

With substitution rate *r*, the codon transition probability matrix *P_ij_*(*t, r*) after time *t* are obtained using matrix exponentiation as

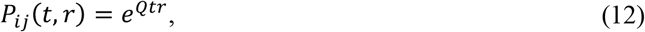

where CSUBST uses the site-wise substitution rate *r_l_* pre-estimated by IQ-TREE and rescaled branch lengths 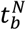 and 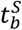 in place of *r* and *t*, respectively. The distribution of expected substitutions at site *l* in branch *k* connecting ancestral node *n* with codon state *i* and a descendant node is therefore given by

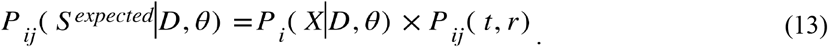

Using *P_klgij_*(*S^expected^*|*D, θ*) in place of *P_klgij_*(*S|D*, *θ*), the total probabilities of expected substitution pairs across sites in the branch pair denoted by *E_C_* are obtained by the same procedure used to obtain *O_C_* (Equation 3 to Equation 8). Similar to *O_C_*, the expected numbers of combinatorial substitutions (*E_C_*) are separately calculated for nonsynonymous and synonymous substitution pairs (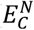 and 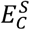, respectively). By definition (Fig. S1C), the following relationships hold at *K* = 2:

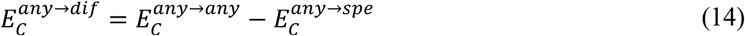

and

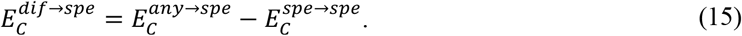

### Nonsynonymous and synonymous combinatorial substitution rates

With the observed and expected numbers of combinatorial substitutions (*O_C_* and *E_C_*, respectively), the rates of nonsynonymous and synonymous combinatorial substitutions are obtained, respectively, by

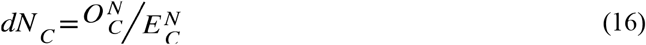

and

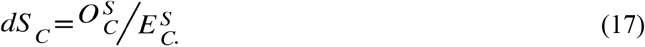

*dN_C_* can be regarded as equivalent to *R* with the per-gene equilibrium amino acid frequencies (their f_gene_), but note that some features are different from the corresponding parts for *R* (Zou and Zhang, 2015a). In particular, we used the standard procedure to derive codon transition probabilities (Equation 13) (Equation 1.2 in (Yang, 2006)), whereas no matrix exponentiation is applied for *R*. In the 21-vertebrate genome dataset, the total expected convergence 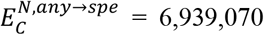 corresponds to 87.2% of the total observed convergence 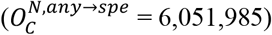. This expectation matches the observation with better accuracy than the previously published results with the *Drosophila* genomes (582.8/932 = 62.5% with their JTT-f_gene_ model) (Zou and Zhang, 2015a).

### Accounting for different range distributions of nonsynonymous and synonymous rates of combinatorial substitutions

Under purifying selection, which is the default evolutionary mode of many proteins (Bustamante et al., 2005), the rate of synonymous substitutions is faster than that of nonsynonymous substitutions. Therefore, saturation of synonymous substitutions becomes a potential problem, especially in a counting method that cannot properly account for the effects of multiple substitutions. To account for this issue, we applied a transformation using quantile values (*U_p_*) as follows:

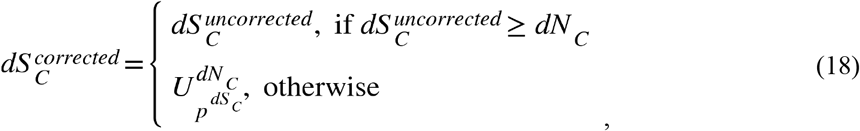

where 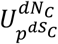 denotes the quantile value of the empirical *dN_C_* distribution at *p^dS_C_^*, the quantile rank of the *dS_C_* value, among all branch combinations. This operation rescales *dS_C_* to match its distribution range with that of *dN_C_*, and the resulting *ω_C_* becomes robust for outlier values (Fig. S13). Because of the need for quantile values, this transformation is only applicable when the branch combinations are exhaustively searched. In this work, 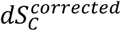 is used at *K* = 2 unless otherwise mentioned.

### Nonsynonymous per synonymous combinatorial substitution rate ratio

A nonsynonymous per synonymous combinatorial substitution rate ratio for *K* branches is given by

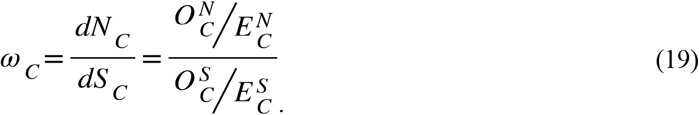

*ω_C_* can be separately calculated for different categories of combinatorial substitutions, e.g., 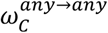 for paired substitutions, 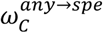 for double divergence, 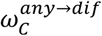 for convergence, 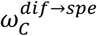 for discordant convergence, and 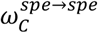 for concordant convergence. For simplicity, the derivation of *ω_C_* was explained above for the combinatorial substitutions illustrated in Fig. S1C. However, our method can be applied to other categories of combinatorial substitutions as well. For example, phenotypic convergence may be associated with the same ancestral amino acid substituted to different amino acids (Konečná et al., 2021), in which case 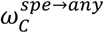 may be useful for analysis.

### Branch combinations

Combinatorial substitutions are a collection of independently occurring evolutionary events (Fig. S1C). Branch combinations containing an ancestor-descendant relationship did not satisfy the evolutionary independence and were therefore excluded from the analysis. Although convergent substitutions occurring in sister branch pairs satisfy the evolutionary independence, they are difficult to discriminate and are often treated as a single ancestral substitution. For this reason, sister branches were also excluded from the analysis (Fig. S10A).

### A branch-and-bound algorithm for the higher-order signature of combinatorial substitutions

*O_C_* and *E_C_*, and hence *ω_C_*, can also be obtained for combinations of more than two branches (*K* > 2). The higher-order analysis is particularly useful when analyzing traits with extensively repetitive convergence, such as C_4_ photosynthesis, which is thought to have evolved at least 62 times independently (Sage et al., 2011). To efficiently explore the higher-order dimensions of branch combinations, we devised a branch-and-bound algorithm that combines the convergence metric cutoff, and the generation of *K* + 1 branch combinations from the branch overlaps at *K* – 1 (Fig. 4A and Fig. S10A). The higher-order analysis starts with an exhaustive comparison of branch pairs (i.e., *K* = 2). Next, convergent branch pairs are extracted with an *ω_C_* cutoff value (≥5.0 in Fig. 4). At this time, branch pairs with a small number of convergent substitutions are excluded by applying an 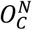 cutoff value (≥2.0 in Fig. 4). The convergent branch pairs are then subjected to the all-vs-all comparison. When a shared branch is found, their union is generated as a combination of three branches to be analyzed. Before proceeding to the analysis at *K* = 3, branch combinations containing a sister or ancestor-descendant relationship are discarded. In this way, *K* is sequentially increased by one at a time. As such, the algorithm searches only for higher-order branch combinations that are guaranteed to have sufficient convergence metrics in lower-order combinations. In each round, convergent branch combinations are first extracted by the cutoffs, and then the *K* + 1 combinations are generated by the *K* – 1 overlap, as in the analysis at *K* = 2. For example, two, three, and four branches should be shared at *K* = 3, *K* = 4, and *K* = 5, respectively. The increase in *K* continues until the algorithm no longer finds a branch combination that satisfies the criteria of *ω_C_* and 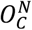.

### Implementation of CSUBST

The proposed methods, including the calculation of *ω_C_* and the branch-and-bound algorithm for higher-order combinations, were implemented in the ‘analyze’ function of CSUBST, which was written in Python 3 (https://www.python.org/). Phylogenetic tree processing was implemented with the python package ETE 3 (Huerta-Cepas et al., 2016). Numpy (Harris et al., 2020), SciPy (Virtanen et al., 2020), and pandas (https://pandas.pydata.org/) were used for array and table data processing. Parallel computation was performed by multiprocessing with Joblib (https://joblib.readthedocs.io/en/latest/). The intensive calculation was optimized with Cython (Behnel et al., 2011).

### Mapping combinatorial substitutions to protein structures

For the analysis of protein structures, a streamlined pipeline was implemented in the ‘site’ function of CSUBST. Using the ‘--pdb besthit’ option, CSUBST requests an online MMseqs2 search (Steinegger and Söding, 2017) against the RSCB Protein Data Bank (PDB) (Berman et al., 2000) to obtain three-dimensional conformation data of closely related proteins. If no hit is obtained, a BLASTP search against the UniProt database is run on the QBLAST server to identify the best-hit protein for which AlphaFold-predicted structure is available (Varadi et al., 2022; Jumper et al., 2021). For some proteins, structural data were manually selected because more appropriate structures were available (e.g., with substrate). Subsequently, CSUBST internally uses MAFFT to generate protein alignments to determine the homologous positions of amino acids and write a PyMOL session file. The protein structures were visualized using Open-Source PyMOL v2.4.0 (https://github.com/schrodinger/pymol-open-source).

### Data visualization

Phylogenetic trees were visualized using the python package ETE 3 (Huerta-Cepas et al., 2016) and the R package ggtree (Yu et al., 2017). General data visualization was performed with python packages matplotlib (Hunter, 2007) and seaborn (Waskom, 2021) as well as the R package ggplot2 (Wickham, 2009, 2). Boxplot elements of all figures are defined as follows: center line, median; box limits, upper and lower quartiles; whiskers, 1.5 × interquartile range.

## Data availability

Raw data and results are available in the Supplementary Dataset (https://doi.org/10.5061/dryad.tx95x6b0v).

## Code availability

CSUBST is available from GitHub (https://github.com/kfuku52/csubst). The results reported in this study can be reproduced with CSUBST v.0.20.17. Scripts used in this study are available in the Supplementary Dataset (https://doi.org/10.5061/dryad.tx95x6b0v).

## Acknowledgments

We acknowledge the following sources for funding: MEXT/JSPS KAKENHI 18J00178 (K.F.), the Sofja Kovalevskaja programme of the Alexander von Humboldt Foundation (K.F.), a Human Frontier Science Program (HFSP) Young Investigators Grant RGY0082/2021 (K.F.), and NIH R01 GM083127 (D.D.P.). Computations were partially performed on the National Institute of Genetics (NIG) supercomputer.

## Author Contributions

K.F. designed the study. K.F. designed and wrote all programs and performed data analysis. D.P. contributed to conceptualizing and helping guide the analysis. K.F. and D.D.P. wrote the paper.

## Supplementary Materials

### Supplementary Texts

**Supplementary Text 1. False positives in the detection of molecular convergence by topology-based methods.** By taking advantage of the branch attraction potentially caused by molecular convergence, which may be detected as a form of site-specific likelihood supports for alternative tree topologies, Parker et al. reported that nearly 200 out of 2,326 orthologous proteins were convergently evolved between echolocating bats and whales (Parker et al., 2013). However, thorough reexaminations of their methodology, which evaluates convergence by phylogenetic tree topology without reconstructing ancestral sequences and substitutions, revealed that most of the reported genomic signatures for molecular convergence were false positives that often lack convergent substitutions (98/117 genes listed as convergent between bats and dolphins), highlighting the need to directly evaluate convergent substitutions rather than indirect signatures such as site-specific likelihood supports (Thomas and Hahn, 2015; Zou and Zhang, 2015b).

**Supplementary Text 2. Phylogenetic combinations of substitutions.** When two separate lineages each experience a codon substitution at the same position in a protein, we call these paired substitutions (Fig. S1C). Paired substitutions may be of interest regardless of the codons involved, particularly if there are coincident bursts of paired substitutions along two lineages and especially if the burst involves more nonsynonymous than synonymous changes. Furthermore, if nonsynonymous paired substitutions result in the same amino acid, they are considered convergent substitutions at the amino acid level, potentially of great interest if similar selective pressures have driven the convergent events. Here, we use the classic definition of convergent evolution, that is when two biological traits in two separate lineages independently evolve to similar endpoints (Pollock and Pollard, 2016). When the paired substitutions in the same codon site result in different amino acids, we call it double divergence or divergent substitutions.

The divergence of the ancestors prior to a convergent event may also be of interest for more complex reasons. First, if the ancestors come from closely related species, the same wild population in the same species, or even replicate populations in the laboratory, the degree of convergence in response to the same selective pressure can be seen as a measure of mechanistic constraint. Convergence under these conditions may indicate that there are only a few easy ways to respond to that selective pressure. At the protein level, amino acid substitutions accumulate combinatorial epistatic effects as they diverge, leading to coevolution (Goldstein and Pollock, 2017). Such coevolution may alter the adaptive landscape but can also lead to decreasing levels of nearly neutral convergence (homoplasy) as proteins diverge. Second, the codon state of the ancestors can strongly affect the accessibility of the convergent state; many types of amino acid substitution are rare in part for this reason, and so convergence events involving one or more rare events may be a stronger indication that they are driven by selection rather than convergence involving common events. We discriminate between two classes of convergent events where the ancestral codon or amino acid states are different (discordant convergence) or the same (congruent convergence). We note that in using this terminology, we are avoiding the term “parallel evolution,” which has rather ambiguous and muddled usage in the literature (Arendt and Reznick, 2008; Pollock and Pollard, 2016) and is sometimes applied to cases of similar or identical ancestral populations, species, biological systems, proteins, or amino acids.

**Supplementary Text 3. New approaches to estimate the rate of molecular convergence.** Among a variety of methods for conventional *ω* estimation (Pond and Frost, 2005; Yang, 2006), the so-called counting methods are most similar to our approach. First, ancestral codon sequences are estimated by the empirical Bayesian method devised in IQ-TREE (Minh et al., 2020), from which the probabilities of codon substitutions are calculated for each branch and site. The substitution probabilities are internally stored in multidimensional arrays designed for efficient processing of substitution probabilities (see Methods). Next, total probabilities of observed combinatorial substitutions (*O_C_*) in a combination of two or more branches are obtained separately for nonsynonymous and synonymous substitutions (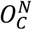 and 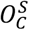, respectively) by deriving joint substitution probabilities with any, different, or specific states at the ancestral and the derived node of a branch (Fig. S1C).

To obtain the total probabilities of expected combinatorial substitutions (*E_C_*), we devised a method that utilizes codon substitution models similar to the previous report that leveraged amino acid substitution models in estimating excess convergence (Zou and Zhang, 2015a) (Fig. S2A). A novel aspect of our approach is that it considers both nonsynonymous and synonymous substitutions. Codon transition probabilities are derived from a mechanistic or empirical codon substitution matrix, empirical codon equilibrium frequencies, branch length, site-wise substitution rates, and the ancestral states of the parent node. Using the expected codon states from this codon transition matrix, the joint probabilities of combinatorial substitutions are calculated as 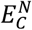 and 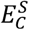, just as in the observed values (see Methods for details).

Finally, after accounting for different ranges of the synonymous and nonsynonymous rates of combinatorial substitutions (*dS_C_* and *dN_C_*, respectively, see Methods for the correction), a formula of the same form as that for calculating conventional *ω* was used to contrast the observed numbers of nonsynonymous and synonymous combinatorial substitutions with their respective expectations to derive *ω_C_* by Equation 19. While *ω_C_* is a general metric that can be calculated individually for different categories of combinatorial substitutions (Fig. S1C), in this work, we consistently discuss the performance of 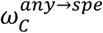, which represents the rate of convergent substitutions, as it is among the most popularly analyzed types of combinatorial substitutions.

Since we will be discussing convergent evolution in the rest of the current study, the superscript *any → spe* will be omitted unless otherwise mentioned.

**Supplementary Text 4. Conventional approaches for estimating convergence rates.** Divergent substitutions have the advantage of being linearly correlated with convergent substitutions (Castoe et al., 2009; Goldstein et al., 2015), although, in *C/D*, the nature of comparing focal branch combinations to the others makes it difficult to identify certain evolutionary scenarios, such as widespread adaptive molecular convergence throughout the tree (Zou and Zhang, 2015a). Expected numbers of convergent substitutions can be obtained from amino acid substitution models (Zhang and Kumar, 1997; Zou and Zhang, 2015a), such as the JTT model (Jones et al., 1992), in combination with observed amino acid frequencies in a protein, an amino acid site, or a group of amino acid sites categorized by the CAT model (Lartillot and Philippe, 2004). However, the difficulty in estimating equilibrium amino acid frequencies from a small number of proteins, especially when per-site frequencies are analyzed, hampers accurate expectations of convergent substitutions (Zou and Zhang, 2015a).

Both methods (utilizing divergent substitutions or expected convergence) successfully recover the pattern of diminishing convergence over time, a recently established evolutionary hallmark of proteins that evolve in the context of intramolecular epistasis (Goldstein et al., 2015; Zou and Zhang, 2015a, 2017). However, false positives are difficult to eliminate due to errors in gene tree topologies caused by technical and biological factors, including incomplete lineage sorting, introgression, and within-locus recombination (Mendes et al., 2016, 2019). Regardless of whether the species tree or individual gene trees are employed, this problem persists as a major source of false convergence in the analysis of genome-scale data.

**Supplementary Text 5. Further evaluations of convergence metrics by simulations.** To further check the robustness of *ω_C_*, we analyzed simulated data under different settings. *ω_C_* was stably estimated under a range of conventional *ω* values (0.1–5.0), indicating that *ω_C_* successfully captures the change in substitution profiles but not the change in the rate of protein evolution (Fig. S3A). A robust estimation was generally achieved even if the codon substitution model was mis-specified in the ancestral reconstruction step (Fig. S3B). One exception was the use of unrealistically simple reconstruction models (MG and GY), in which the variances of *dN_C_* and *ω_C_* increased while the median did not change greatly. Therefore, care should be taken when a simple model is used. *ω_C_* was robust against other factors, as mentioned in the main text (Fig. S3C–G).

**Supplementary Text 6. Signature of intramolecular epistasis in empirical convergence.** In the known examples of adaptive protein convergence, we found that the rate of concordant convergence 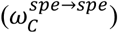 is significantly higher than that of discordant convergence 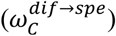, with the largest contribution to the *χ*^2^ statistic coming from depleted nonsynonymous substitutions in discordant convergence (Fig. S4J–K, *P*-value is shown in the plot). Such a pattern was not detected in the simulated adaptive convergence (Fig. S3A). The simulated codon sequence evolution assumes independence between sites; therefore, intramolecular epistasis is ignored. In the presence of epistasis between amino acid sites, a substitution at one site will change the substitution profiles of other coupled sites (Pollock et al., 2012), and subsequent substitutions in the coupled sites entrench the original site (Goldstein and Pollock, 2017; Shah et al., 2015; Starr et al., 2018). This means that epistasis makes it difficult to replace different ancestral amino acids with the same derived amino acid, even in homologous sites in the same protein (Fig. S4L). Thus, intramolecular epistasis can be a source of the different rates between concordant and discordant convergence.

**Supplementary Text 7. Decreasing rates of combinatorial substitutions over time.** To further characterize rate decreases over time, we took advantage of the ability to apply *ω_C_* to a variety of combinatorial substitutions. We asked whether the rate decrease is specific to convergence by performing the same analysis for other categories of combinatorial substitutions (Fig. S1C). Notably, the rate of double divergence decreased over time in a manner similar to the decrease in convergence (Fig. S7B). The sum of double divergence and convergence corresponds to paired substitutions (Fig. S1C), the rate of which also decreased over time (Fig. S7C). These results suggest two possibilities. One result is that epistatic changes from neighboring amino acid residues impose constraints on not only to which amino acid state a site tends to substitute (i.e., site-specific substitution profile), but also on which amino acid sites tend to substitute (i.e., site-specific substitution rate). The alternative (not necessarily exclusive) possibility is that doubly divergent events are decreasing because the rate of convergence to similar but not identical amino acids decreases just as the rate of convergence to identical amino acids decreases. In either case, this effect may be important to account for in analyses of adaptation.

**Supplementary Text 8. Potential artifacts by false gene grouping.** We sometimes observed anomalously high synonymous convergence rate (*dS_C_*) in extremely distant branch pairs, which can be attributed to an incorrect grouping of different gene families. Although orthogroup inference has dramatically improved in accuracy in recent years (Emms and Kelly, 2015, 2019), it does not completely eliminate false groupings. In line with this idea, orthogroups that encompass extremely large genetic distances tend to contain multiple sets of genes that have clearly non-homologous sets of protein domains (Fig. S7E; Supplementary Dataset for orthogroups with total branch distance greater than 15 nucleotide substitutions per nucleotide site). In any case, such artifacts were successfully captured by *dS_C_* and corrected for in *ω_C_*.

**Supplementary Text 9. Genome editing as a means to evaluate the mutational effects of molecular convergence.** The rapid development of genome editing technologies with CRISPR/Cas-based systems (Anzalone et al., 2020; Knott and Doudna, 2018) provides a means to test the effect of mutations on *in vivo* phenotypes using targeted mutagenesis. This approach can help us understand important biological processes, for example, for livestock and crop enhancements. However, because of the massive mutations accumulated in the lineage of interest, a key challenge is the efficient identification of important mutations, and even more so for combinations of mutations because mutational effects are often dependent on genetic background (Chandler et al., 2013). Convergent evolution, which can be seen as replicated experiments by nature, has the potential to solve this problem. Convergent mutations that arise in different lineages are likely to have stronger effects and depend less on the genetic background than mutations that were not convergent under the same physiological or phenotypic adaptive pressure, and such mutations and the genes that carry them are thus promising candidates to achieve desired phenotypes. One successful example is the toxin resistance conferred to an engineered fruit fly strain, “monarch fly,” which harbors convergent amino acid substitutions, also found in monarch butterflies, in its sodium pump ATPalpha1 (Karageorgi et al., 2019; Taverner et al., 2019). As such, adaptive molecular convergence discovered by our method could be experimentally verified while utilizing genome editing.

**Supplementary Text 10. Protein size–dependent change in convergence rates.** The genome-scale analysis of vertebrate genes allowed us to correlate various protein properties with convergence rates. In the course of analysis, we found that protein sizes negatively correlate with convergence rates (ρ = −0.11 with *C/D* and ρ = −0.11 with *dN_C_*; Fig. S11A). Unlike the temporal variation, it is difficult to explain this trend with epistasis because larger proteins should have more epistatic interactions that increase convergence probability (Goldstein et al., 2015; Lyons et al., 2020; Zou and Zhang, 2015a). In addition, protein size does not correlate with genetic distance (ρ = 0.01; Fig. S11B), confirming that confounding is negligible. A similar trend in synonymous convergence rate (ρ = −0.07 with *dS_C_*) suggests that, unlike the temporal variation (Fig. 2B), the pattern is largely nonbiological and perhaps created by the uncertainty caused by the small number of codon substitutions in small genes. As the trend is consistently observed in nonsynonymous and synonymous convergence rates, *ω_C_* was relatively stable over protein size (ρ = −0.06), further demonstrating its robustness against artifacts.

**Supplementary Text 11. Remarks on empirical datasets.** For benchmarking, we collected known examples of molecular convergence associated with phenotypes. While we followed the same taxon sampling as in the original reports (cited in the main text), further additions and scrutiny of taxons allowed us to find previously unappreciated features in some datasets.

The convergence of mitochondrial proteins between snakes and lizards of the Agamidae family was reported previously (Castoe et al., 2009). In our mitochondrial genome dataset, a massive burst of amino acid convergence was found between snakes and Acrodonta, the lineage consisting of not only Agamidae but also Chamaeleonidae. This detail was not in the previous report because Chamaeleonidae were not available at the time to be included in the phylogenetic analysis.

Improved phylogenetic resolution is known to increase the specificity of convergent site detection (Thomas et al., 2017). In carnivorous plants, several amino acid substitutions were reported previously in digestive enzymes (Fukushima et al., 2017). With additional plant genomes (Table S8), the candidate convergent substitutions were narrowed down in this study to smaller numbers of substitutions that correlated more tightly in the phylogenetic placement with the evolution of carnivory. One of the convergent substitutions found in both the previous report and this study is located at a substrate-binding site in the family GH19 chitinases (Fig. S4F). Double divergence was found in a substrate-binding site of PAPs (Fig. S4G).

**Supplementary Text 12. Use of posterior probabilities of ancestral states for the inference of substitutions.** To estimate the posterior probabilities of substitutions, we sum over the posterior probabilities of ancestral states. In this way, we circumvent a computationally expensive step employed in previous reports to handle individual Markov chain Monte Carlo (MCMC) samples separately (Fukushima et al., 2017; Goldstein et al., 2015). However, since the posterior probabilities are not independent for each node of a phylogenetic tree, this approximation comes at the expense of accuracy in estimating substitution probabilities. In the analysis of amino acid sequences, it is difficult to exclude such a bias. In contrast, in our method, this bias appears in both nonsynonymous and synonymous substitutions and is likely to be canceled out when calculating *ω_C_*, the ratio of their convergence rates. To assess the impact of summing over the ancestral state posteriors, we reanalyzed the vertebrate genome dataset with the CSUBST option --ml_anc to binarize the posterior probabilities in the three-dimensional arrays with the size of *M* × *L* × 61 (see Methods). This operation corresponds to the uniformization between MCMC samples, and the substitution probabilities are binarized accordingly. In this setting, we reproduced the analysis shown in Fig. 2B. Although the temporal trends were consistent, the convergence metrics, especially *dN_C_* and *dS_C_*, were slightly higher than those in Fig. 2B (i.e., more conservative without binarization) (Fig. S12). Importantly, such a shift was less evident in *ω_C_*, as expected. These observations led us to adopt the approximation of substitution probabilities in the *ω_C_* calculation to take advantage of computational speed-up.

### Supplementary Tables

**Table S1. Methods to detect convergent signatures of protein sequences.** (separate file)

**Table S2. Parameter settings for the simulated molecular evolution.** (separate file)

**Table S3. Summary of empirically validated protein convergence.** (separate file)

**Table S4. Convergence statistics in empirically validated protein convergence.** (separate file)

**Table S5. List of branch pairs with herbivory-associated protein convergence.** (separate file)

**Table S6. List of branch pairs where simultaneous convergence of gene expression and protein sequences is detected.** (separate file)

**Table S7. Time required for the analysis of higher-order convergence in PEPC.** (separate file)

**Table S8. Genome and transcriptome data.** (separate file)

### Supplementary Figures

**Figure S1.**
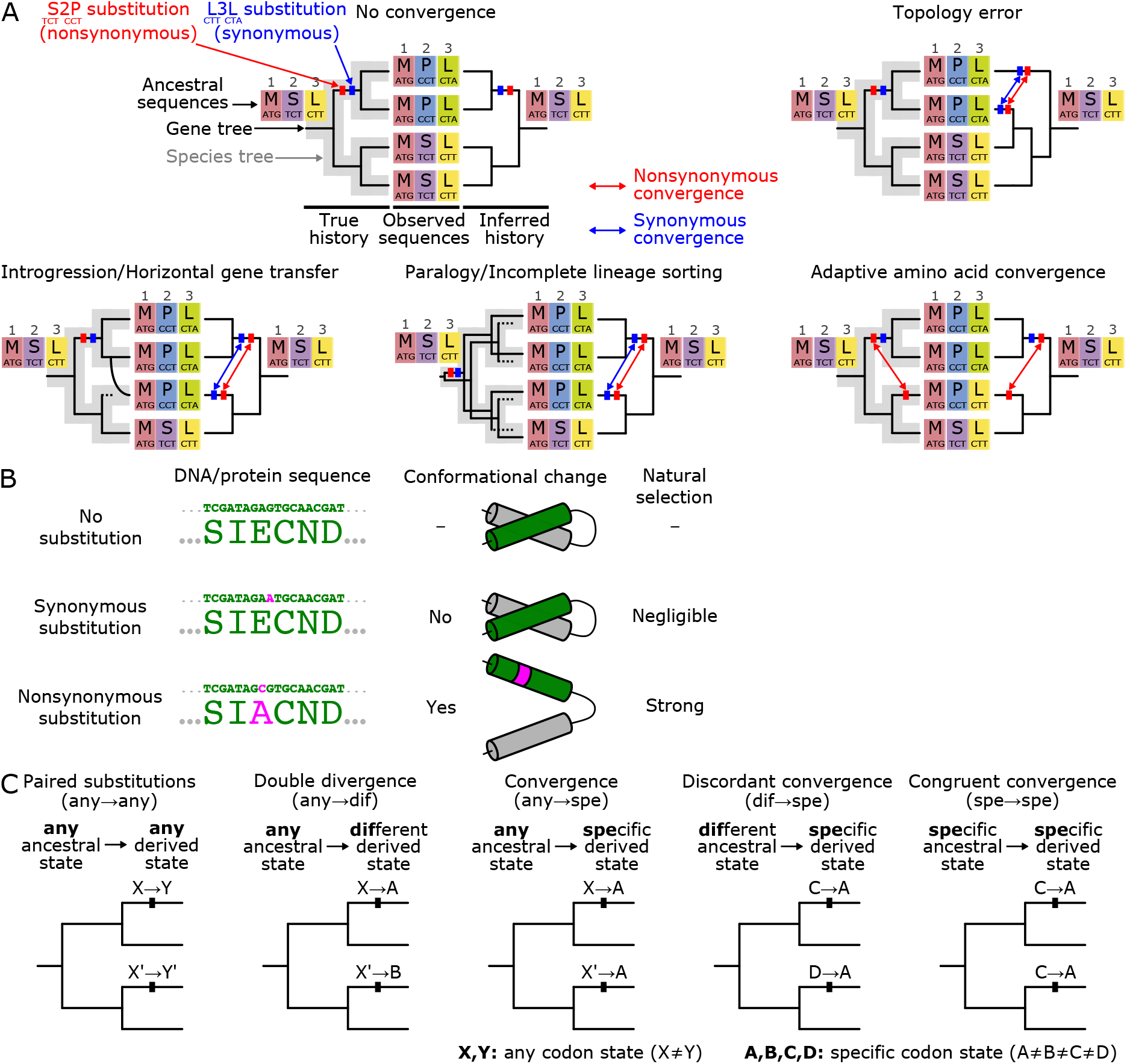
Types of substitution and their relationships to evolutionary patterns. (**A**) Errors in tree topology lead to false convergence. No convergence is detected as long as the phylogenetic tree is correctly inferred, while errors in the tree topology can lead to spurious convergence. Even if the species tree is correctly inferred, there can still be spurious convergence if introgression or horizontal gene transfer (HGT) has occurred. A similar situation can arise from paralogy and incomplete lineage sorting. While the above technical and biological factors alter the inference of both nonsynonymous and synonymous substitutions, adaptive convergence should involve an increased rate of nonsynonymous convergence without changing synonymous convergence. (**B**) The relationship between the type of substitution, protein conformation, and natural selection. (**C**) Combinatorial substitutions with evolutionary importance. A pair of substitutions at the same site in two lineages are annotated on branches (ancestral→derived). X and Y indicate any codon state, and A, B, C, and D denote specific codon states.

**Figure S2.**
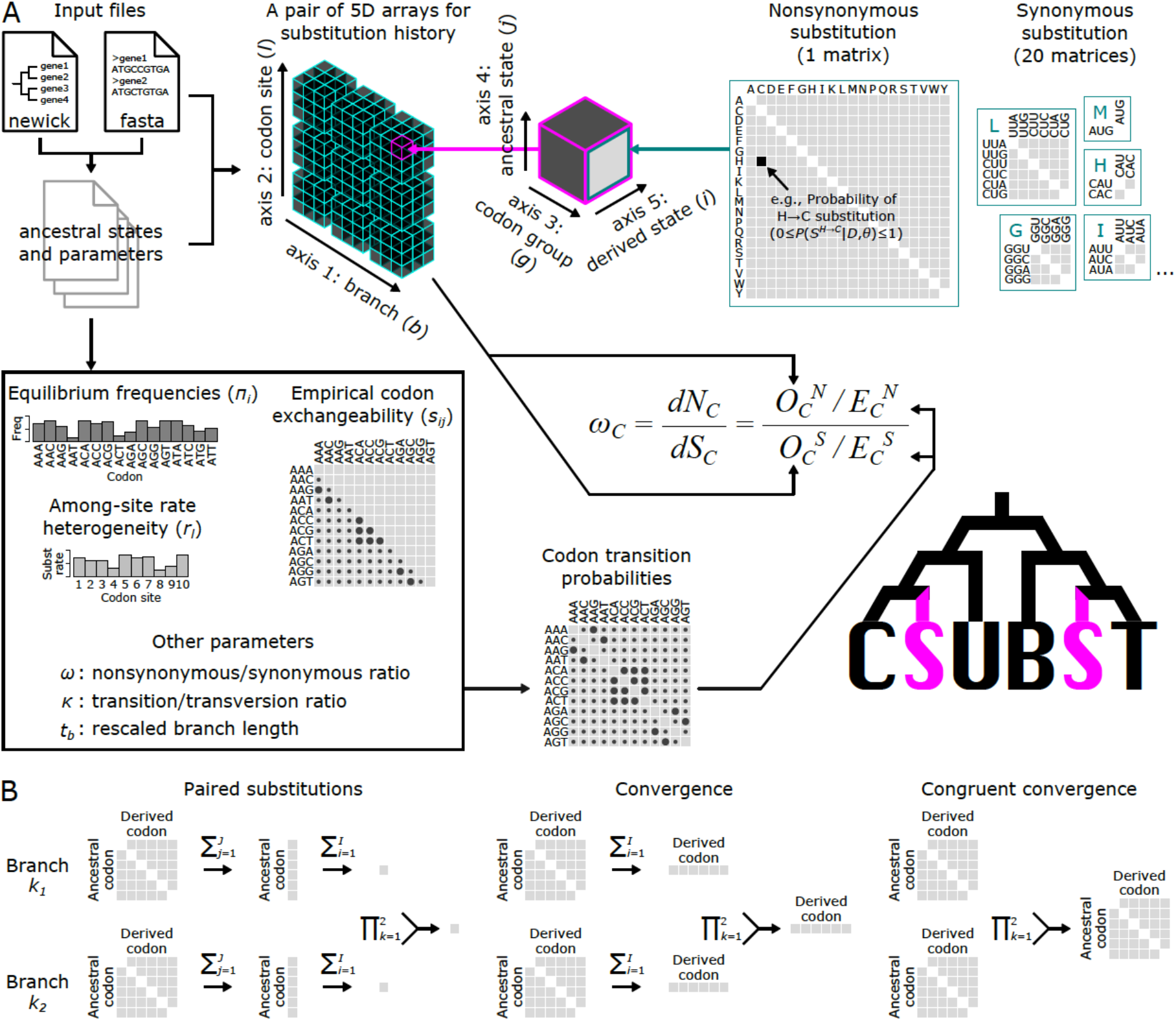
Overview of the method. (**A**) Flow of data in CSUBST. (**B**) Array operations for deriving the probabilities of combinatorial substitutions.

**Figure S3.**
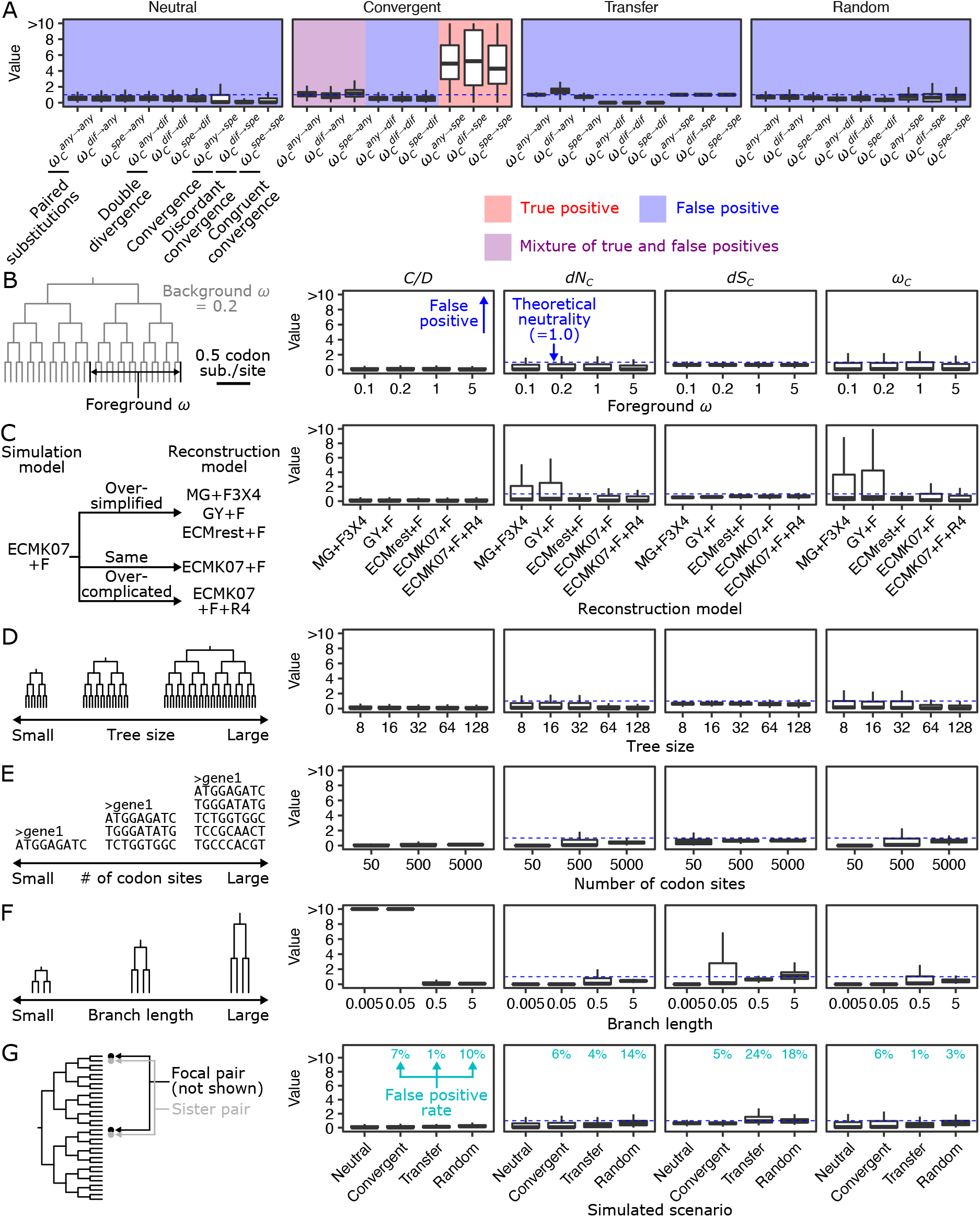
Robustness of convergence metrics under simulated conditions. (**A**) Comparison of the complete set of *ω_C_* variants. There are nine *ω_C_* variants, of which three are associated with convergence: 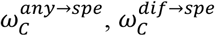 and 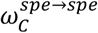. (**B**) Conventional *ω* values. According to the value of *ω*, the mode of protein evolution can be categorized into purifying selection (*ω* < 1), neutral evolution (**ω** = 1), and adaptive evolution (**ω** > 1). The examined parameters are illustrated on the left in **B-G**. If no changes are indicated, the parameters of the simulations are the same as in the “Neutral” scenario in Fig. 1C,D. To the right, each box plot corresponds to the results of 1,000 simulations. Dashed lines indicate the neutral expectation (=1.0) except for *C/D*, for which no theoretical expectation is available. (**C**) Model misspecifications. The following base models were analyzed: MG (Muse and Gaut, 1994), GY (Goldman and Yang, 1994), ECMrest (Kosiol et al., 2007), and ECMK07 (Kosiol et al., 2007). (**D**) Tree sizes. (**E**) Number of codon sites. (**F**) Branch lengths. When the branch length equals 1, an average of one substitution occurs per codon site. (**G**) Sister branches. The pairs of branches sister to focal branches in Fig. 1C,D were analyzed.

**Figure S4.**
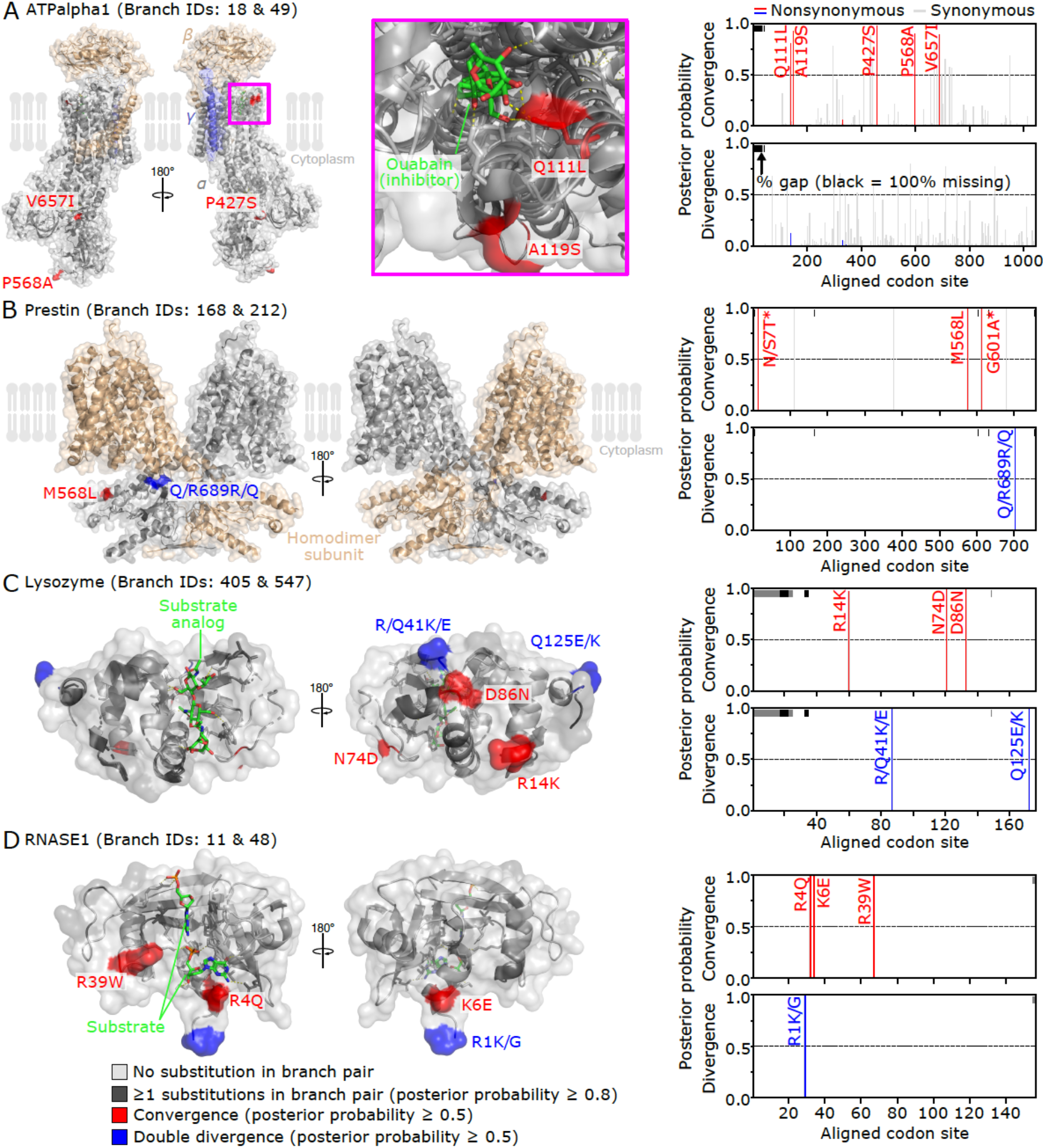
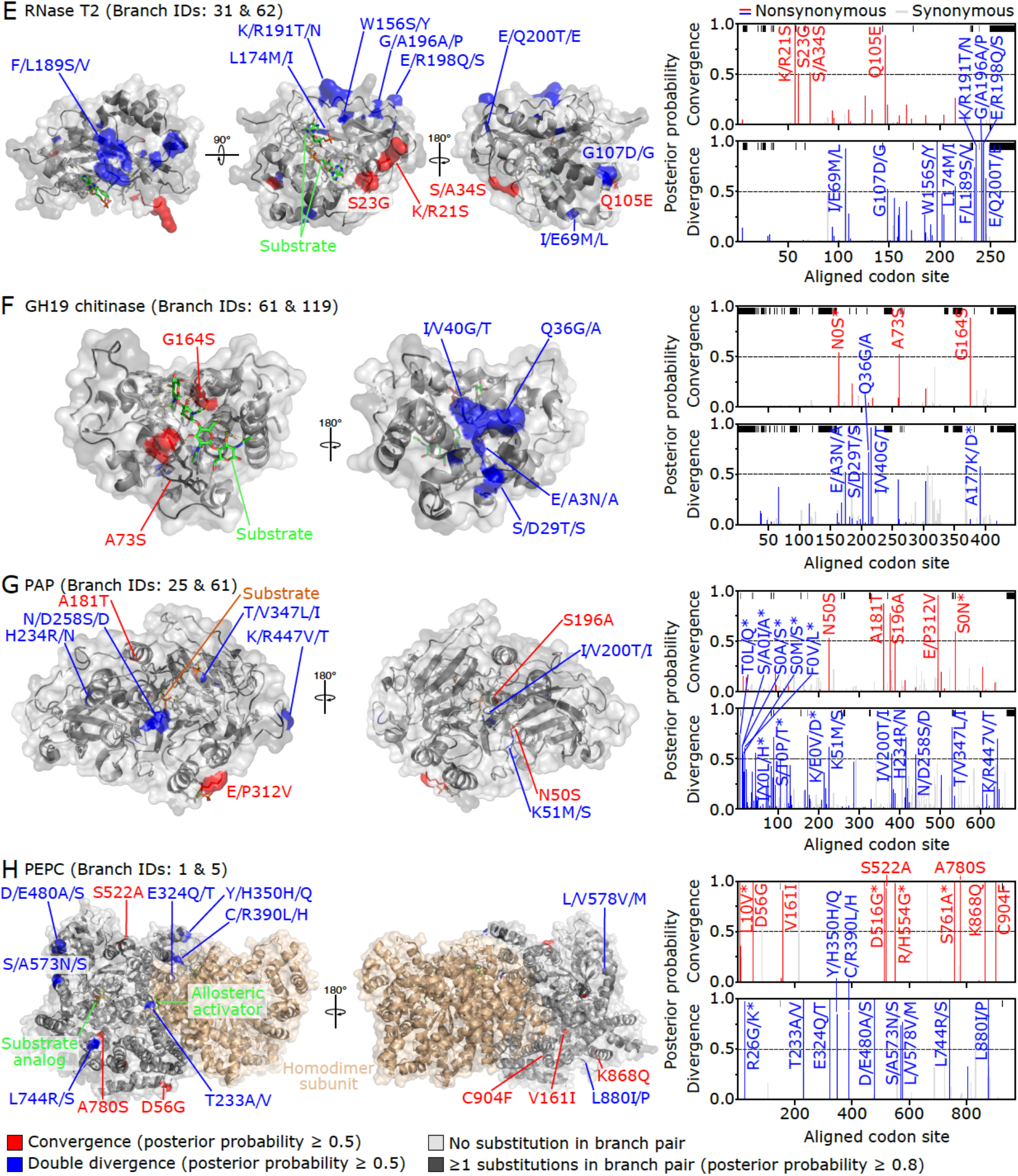
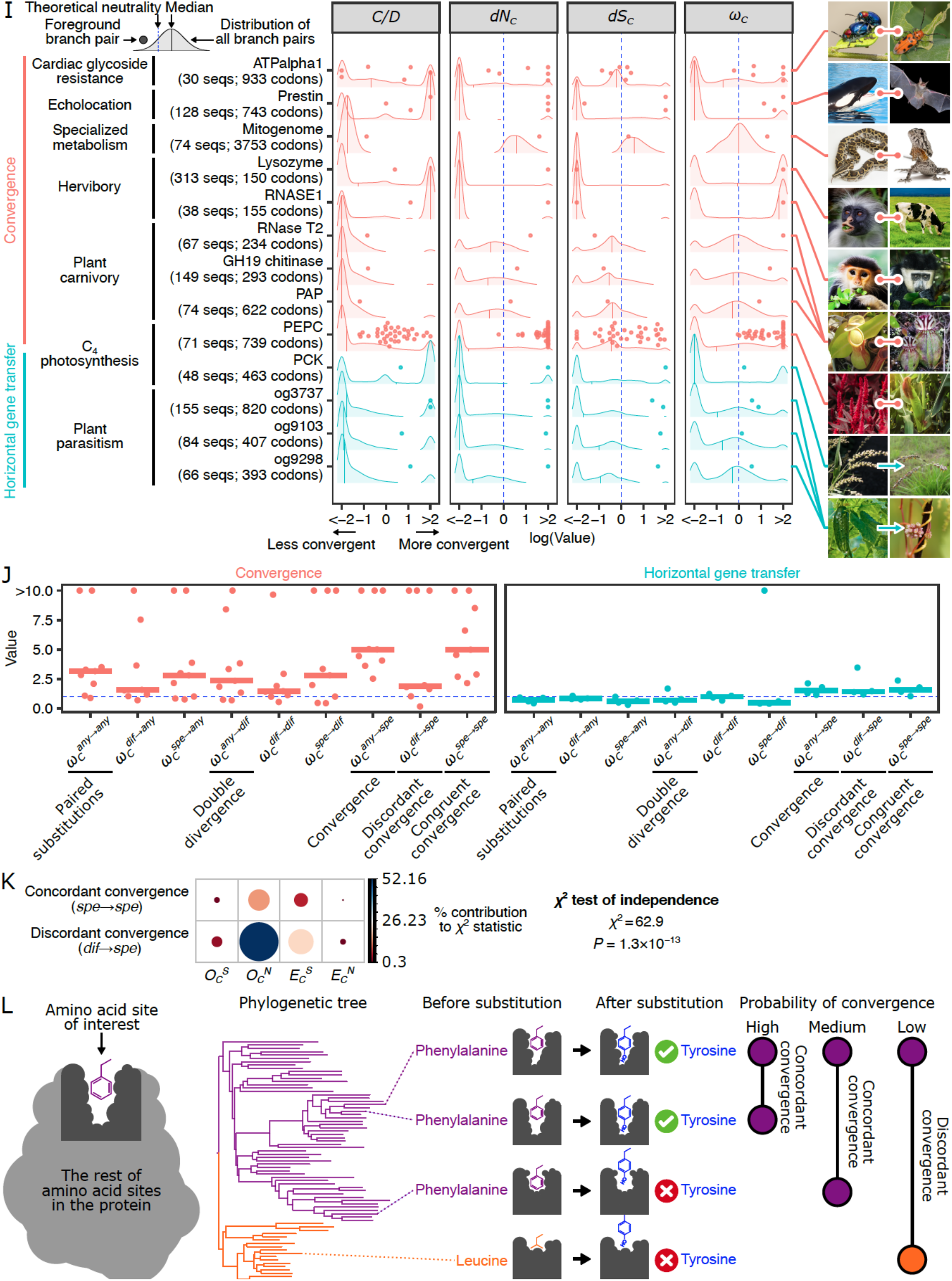
Convergence metrics in genes associated with phenotypic convergence. (**A–H**) Mapping of combinatorial substitutions to the protein structures of ATPalpha1 (**A**, PDB ID: 4HYT), Prestin (**B**, 7LGU), Lysozyme (**C**, 9LYZ), RNASE1 (**D**, 2QCA), RNase T2 (**E**, 1VCZ), GH19 chitinase (**F**, 4IJ4), PAP (**G**, 6GIZ), and PEPC (**H**, 6MGI). The surface representation of the protein is overlaid with a cartoon representation. Convergent and divergent amino acid loci are highlighted in red and blue, respectively. Substrates and their analogs are shown as green sticks. Side chains forming the substrate-binding site are also shown as sticks. Note that these are the side chains in the protein from databases, so amino acid substitutions in the convergent lineages may result in distinct structures and arrangements. The probability of combinatorial substitution for each codon site is shown to the right. Asterisks indicate sites that are not included in the PDB protein structure. Site number 0 indicates no homologous site in the PDB protein structure. A representative branch pair is shown when three or more convergent lineages exist. (**I**) Known examples of protein convergences and HGTs were analyzed with *C/D, dN_C_*, *dS_C_*, and *ω_C_*. Encoded proteins, associated traits, and numbers of sequences and codon sites are provided along the y-axis labels. The images to the right depict the organisms representative of the focal lineages. Points correspond to individual pairs of branches in the gene tree (shown in Fig. S5 and Fig. S6). The photograph of *Alloteropsis semialata* is licensed under CC BY-SA 3.0 (https://creativecommons.org/licenses/by-sa/3.0/) by Marjorie Lundgren. (**J**) Comparison of the complete set of *ω_C_* variants. Points correspond to individual gene trees. Horizontal bars indicate median values. (**K**) *χ*^2^ test comparing the number of combinatorial substitutions associated with concordant convergence and those associated with discordant convergence. The number of combinatorial substitutions in all focal branch pairs of known protein convergence was summed. Circle sizes and colors indicate the relative contribution to the *χ*^2^ statistic. (**L**) Schematic representation of the relationships between intra-molecular epistasis and the rates of convergence. As the inter-branch distance increases, the local environment around the amino acid site changes in the protein structure, leading to a change in the propensity of amino acid substitutions (Goldstein and Pollock, 2017; Goldstein et al., 2015).

**Figure S5.**
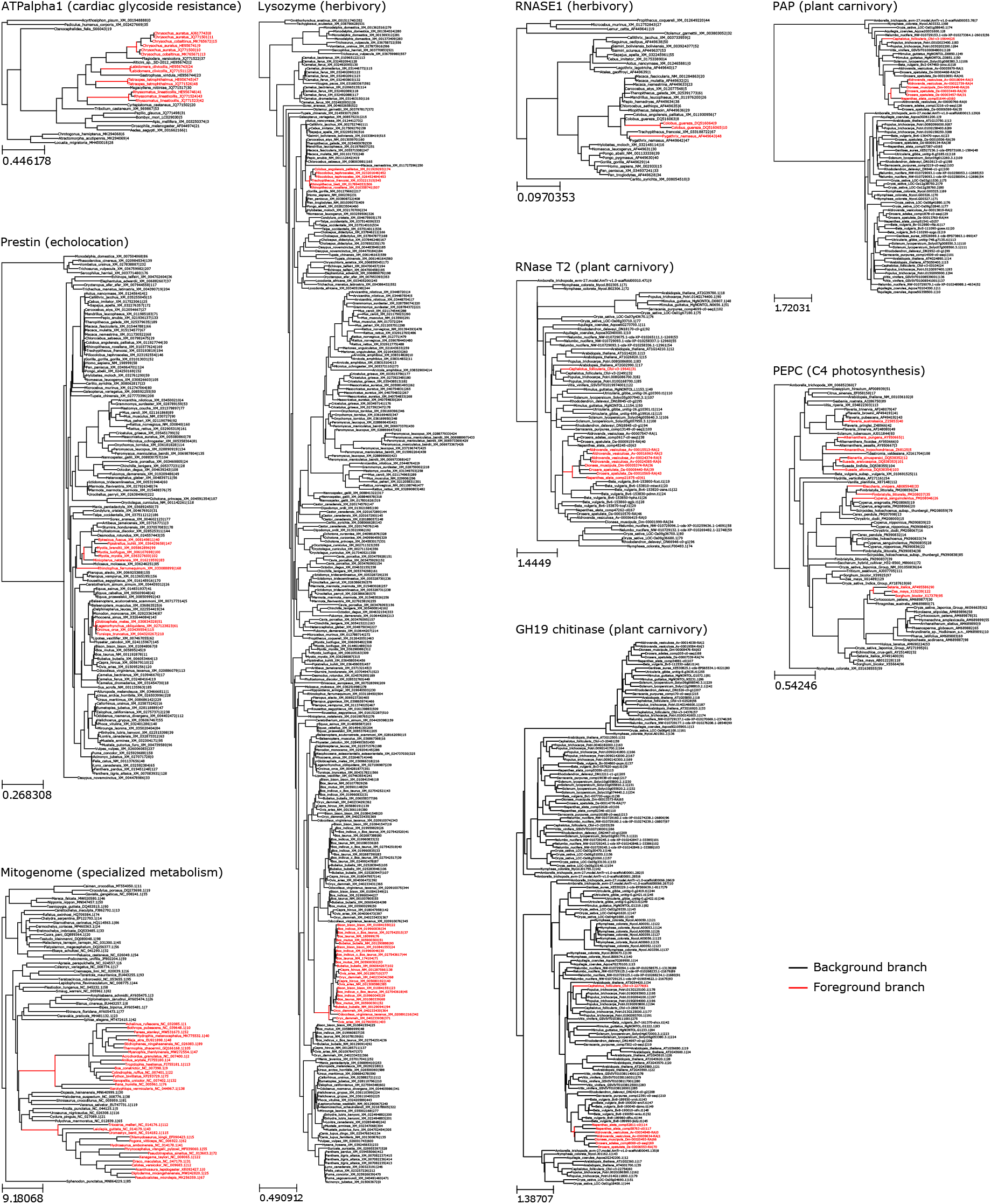
Maximum-likelihood phylogenetic trees for the reported cases of convergent evolution. Scale bars indicate substitutions per nucleotide site. Red indicates focal branches (Fig. 1E).

**Figure S6.**
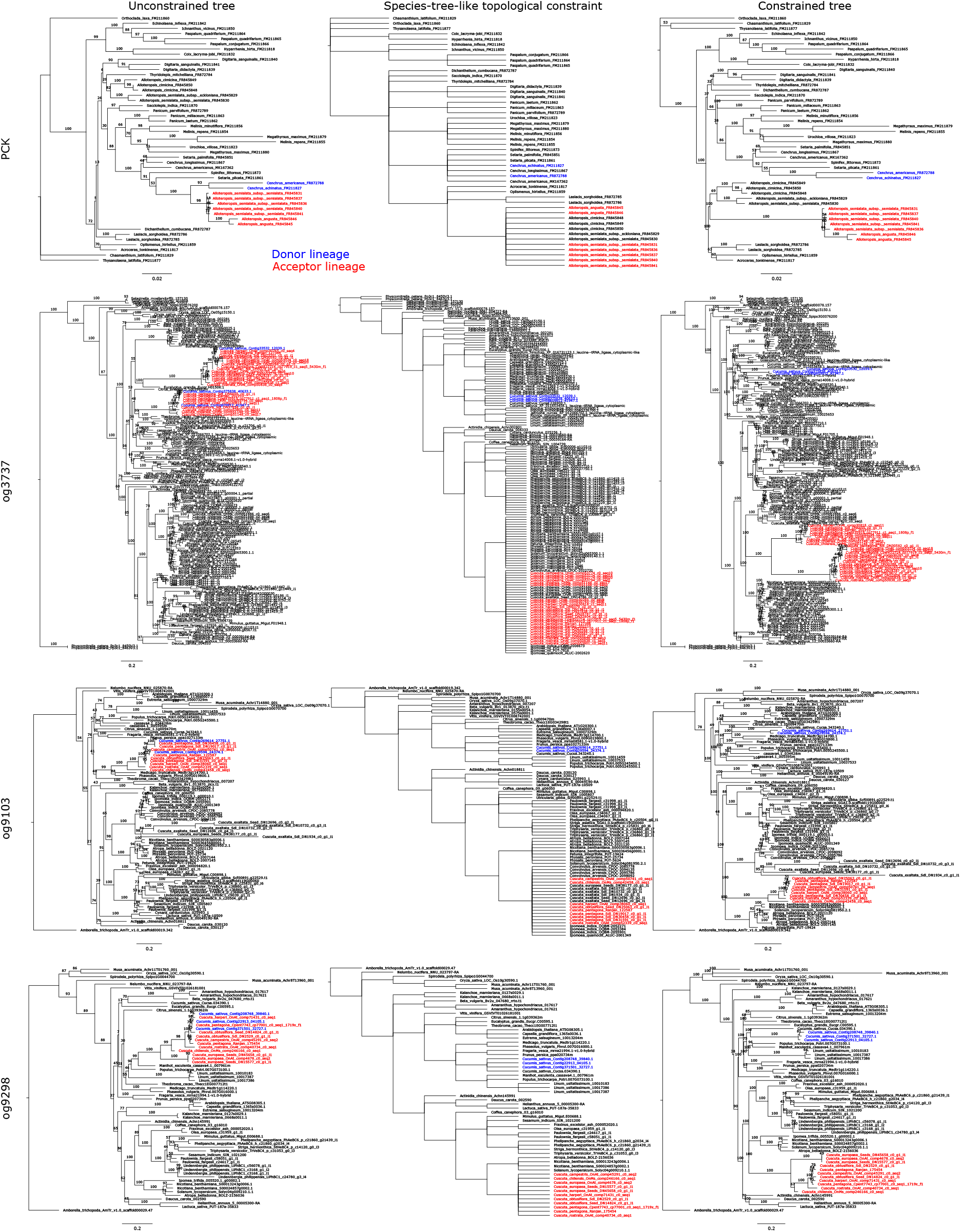
Introducing the species-tree-like topology in the phylogenetic trees involving HGTs. Without a tree constraint, donors and acceptors form a sister clade in the maximum-likelihood phylogenetic analysis (left). When the taxonomic rank information is employed as a constraint in the topology inference (middle), the resulting trees inherit such topologies where donors and acceptors are separated (right). The constrained trees are used to examine how different metrics behave upon false convergence caused by the species-tree-like topology (Fig. 1E). Scale bars indicate substitutions per nucleotide site. Numbers on branches denote ultrafast bootstrapping values (also available as Newick files in Supplementary Dataset).

**Figure S7.**
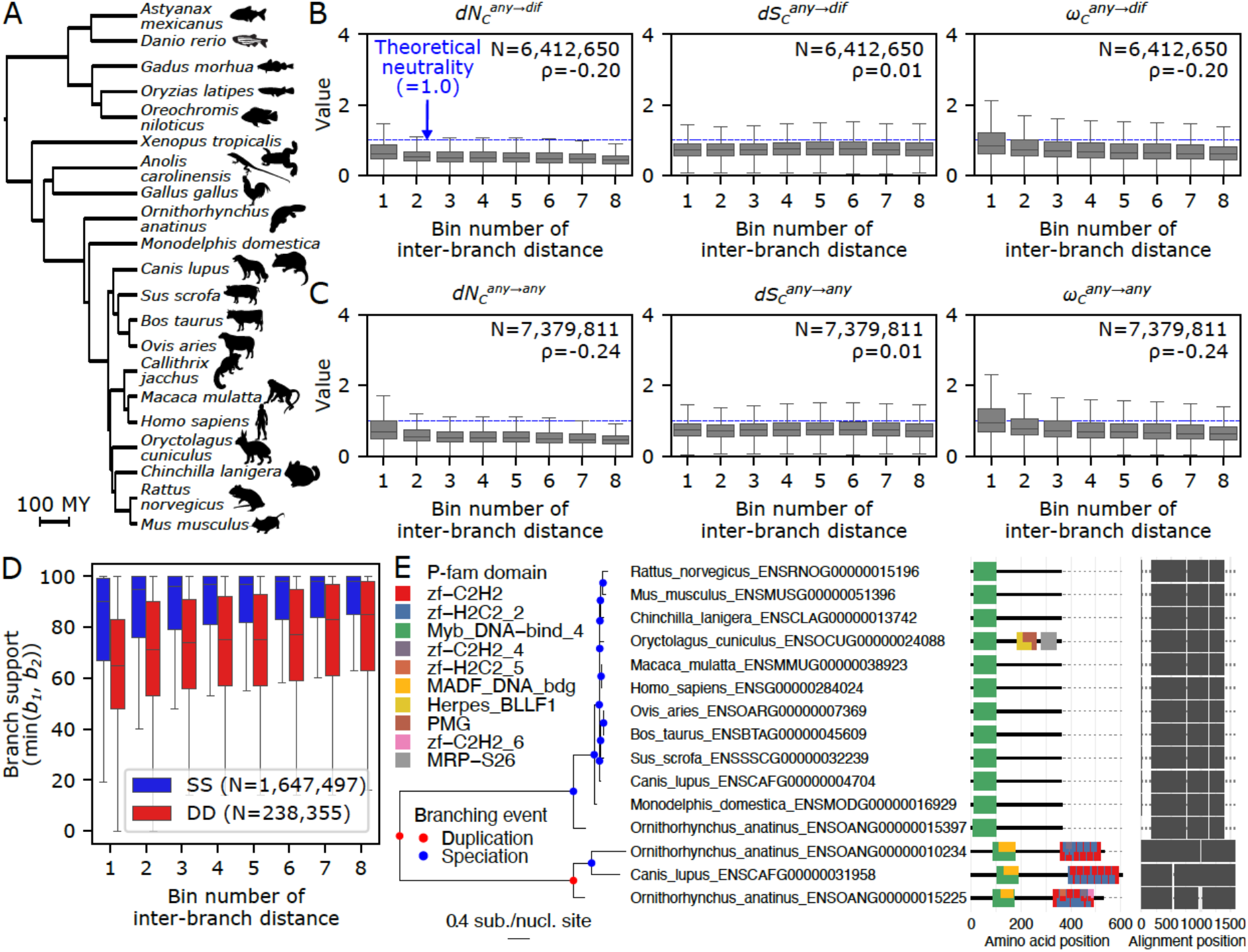
Genome-scale analysis of convergence in nuclear-encoded genes. (**A**) The vertebrate species tree for the 21 analyzed genomes. Some animal silhouettes were obtained from PhyloPic (http://phylopic.org). The silhouettes of *Astyanax mexicanus* and *Oreochromis niloticus* are licensed under CC BY-NC-SA 3.0 (https://creativecommons.org/licenses/by-nc-sa/3.0/) by Milton Tan (reproduced with permission), and those of *Anolis carolinensis* (by Sarah Werning), *Ornithorhynchus anatinus* (by Sarah Werning), and *Rattus norvegicus* (by Rebecca Groom; with modification) are licensed under CC BY 3.0 (https://creativecommons.org/licenses/by/3.0/). (**B**) Temporal variation of double divergence rates. The number of branch pairs (N) and Spearman’s correlation coefficients (ρ) are provided in the plot. (**C**) Temporal variation of paired substitution rates. (**D**) Branch supports in relation to gene duplication. The IQ-TREE’s ultrafast bootstrap values are compared. Reconciled branches were treated as no support (= 0). (**E**) An orthogroup that contains extremely large genetic distances. The gene tree of OG0007724 is shown as an example. Node colors in the trees indicate inferred branching events of speciation (blue) and gene duplication (red). Two clades are connected by an extremely long branch and have non-homologous sets of protein domains. The placement and identity of P-fam protein domains (*E* value < 0.01) are shown to the right of the tree.

**Figure S8.**
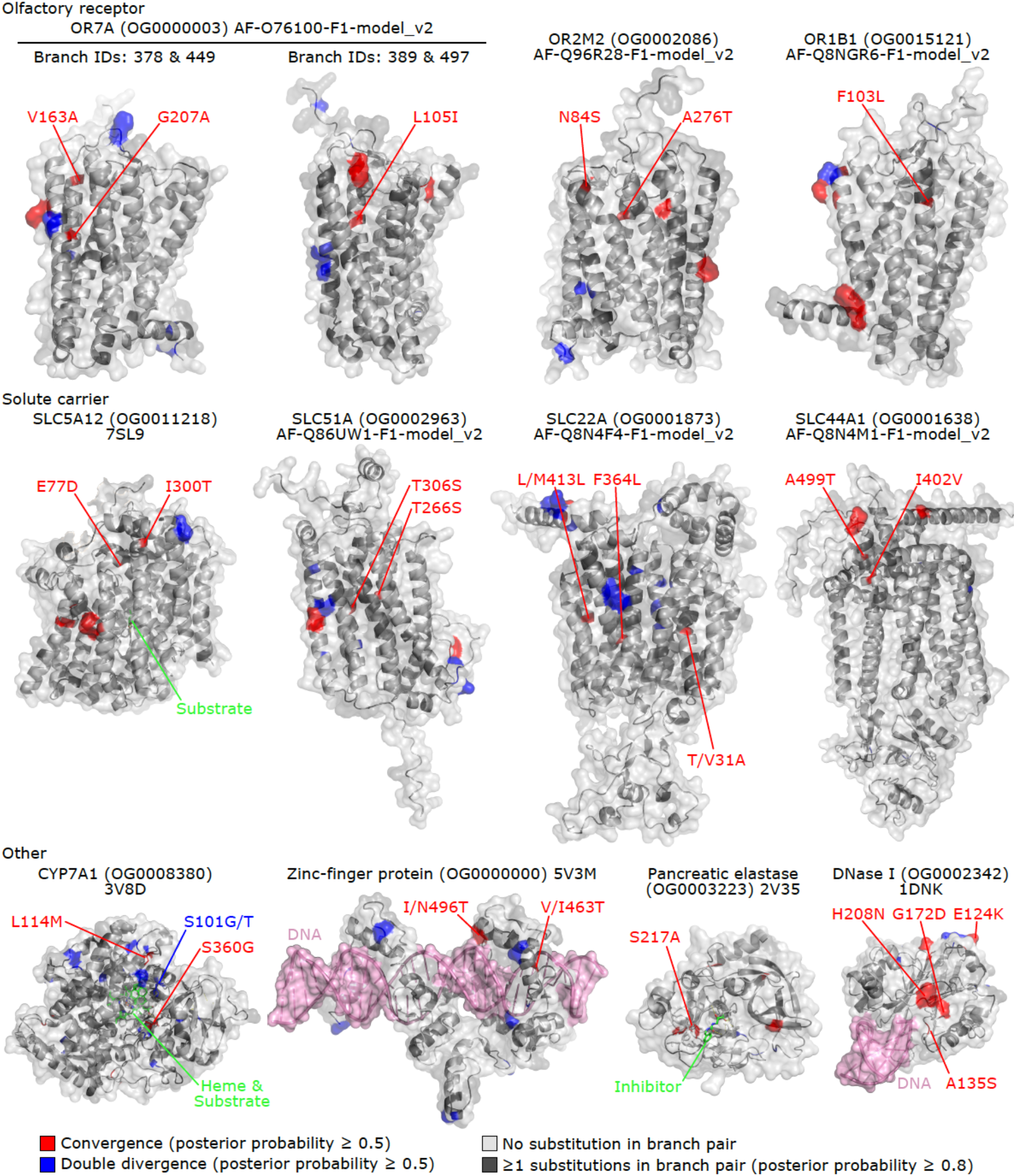
Examples of proteins convergently evolved in herbivores. Convergently evolved proteins (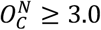 and *ω_C_* ≥ 3.0) in ruminants (*Bos taurus* and *Ovis aries*) and rabbits (*Oryctolagus cuniculus*) are shown (for a complete list, see Table S5). Convergent amino acid substitutions discussed in the main text are labeled. Site numbers correspond to those in the PDB entry or the AlphaFold structure (accession numbers are indicated in the plot). Olfactory receptors and solute carriers are transmembrane proteins, and the upper portion of each protein corresponds to the extracellular region. The surface representation of the protein is overlaid with a cartoon representation. Convergent and divergent amino acid loci are highlighted in red and blue, respectively. Substrates and their analogs are shown as green sticks. Side chains forming the substrate-binding site are also shown as sticks. Note that these are the side chains in the protein from databases, so amino acid substitutions in the convergent lineages may result in distinct structures and arrangements.

**Figure S9.**
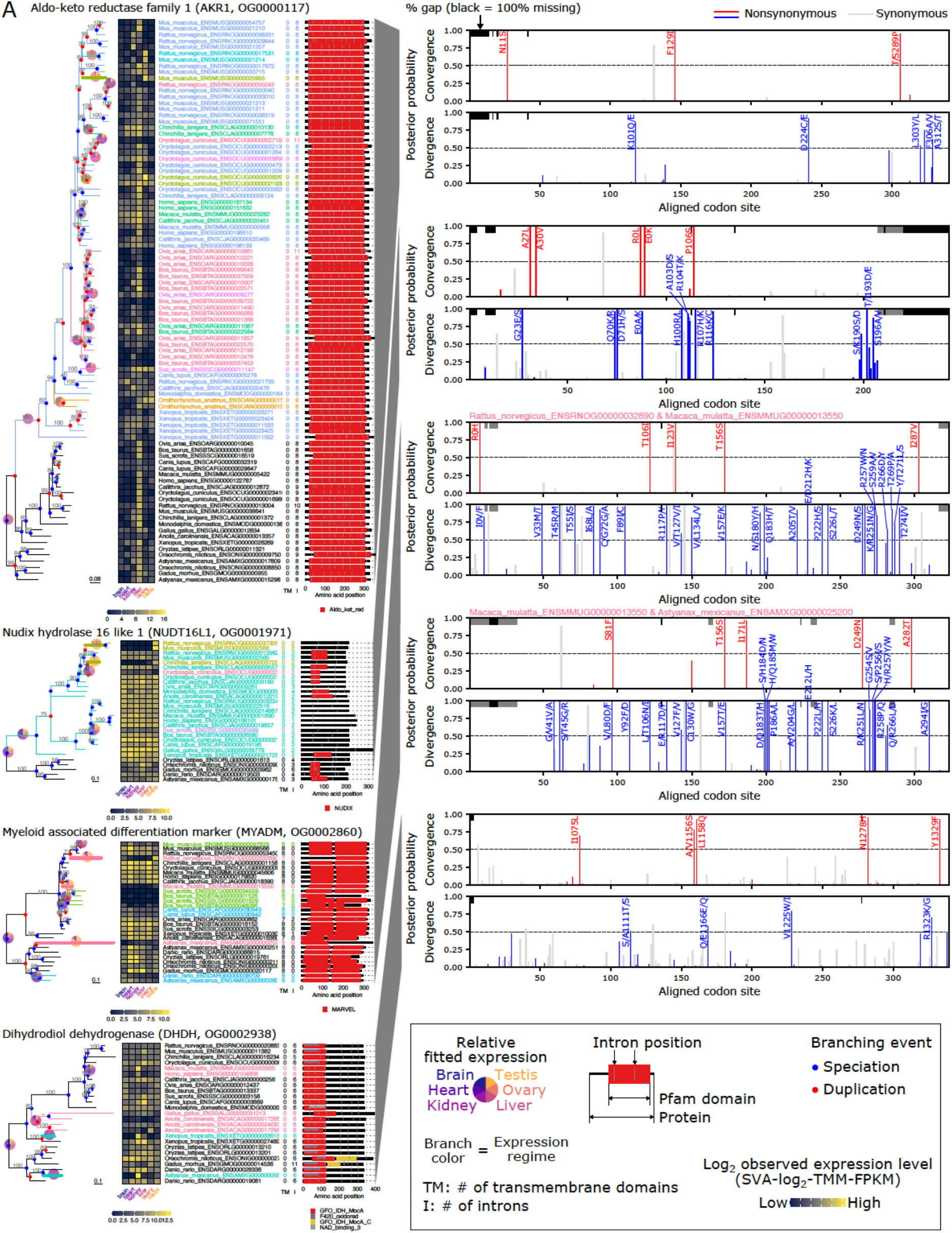
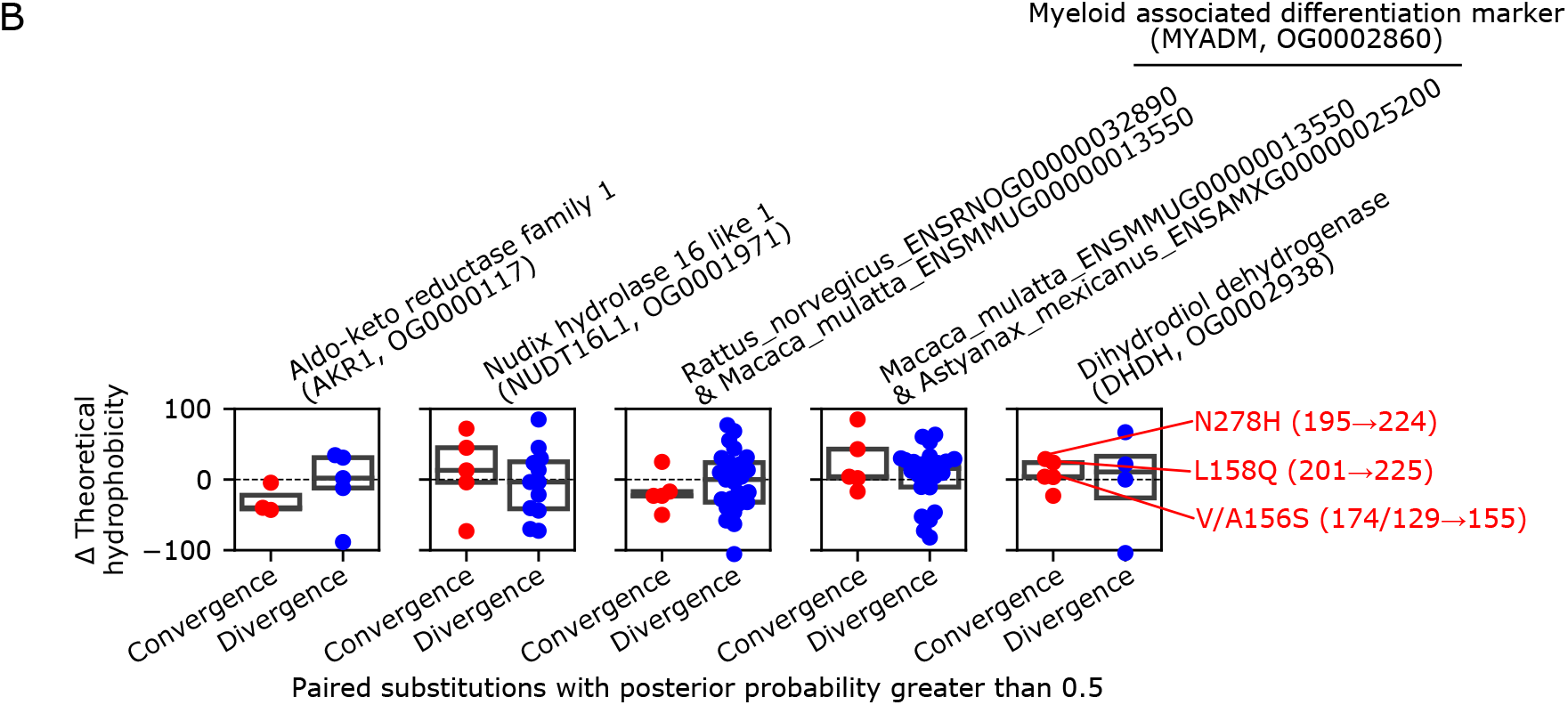
Further characterization of protein convergence jointly occurring with gene expression convergence. (**A**) Complete phylogenetic trees and site-wise posterior probabilities of convergence and divergence in the detected branch pairs. IQ-TREE’s ultrafast bootstrap values are shown above branches. A hyphen (-) marks a branch reconciled by GeneRax. Node colors in the trees indicate inferred branching events of speciation (blue) and gene duplication (red). The heatmap shows expression levels observed in extant species. The colors of branches and tip labels indicate expression regimes. Among-organ expression patterns are shown as a pie chart for each regime. Branches involved in joint convergence are highlighted with thick lines. To the right of the tip labels, the number of transmembrane domains predicted by TMHMM (Krogh et al., 2001), the number of introns in protein-coding sequences, and the Pfam domain structures (E-value < 0.01) are shown. Trees are available as pdf files in Supplementary Dataset. (**B**) Hydrophobicity change of combinatorial amino acid substitutions. Theoretically derived hydrophobicity scales (Tien et al., 2013) were compared between the average values of ancestral and derived amino acids (Δ theoretical hydrophobicity; mean derived amino acid hydrophobicity - mean ancestral amino acid hydrophobicity). Convergent substitutions at the substrate-binding sites of DHDH are labeled and discussed in the main text.

**Figure S10.**
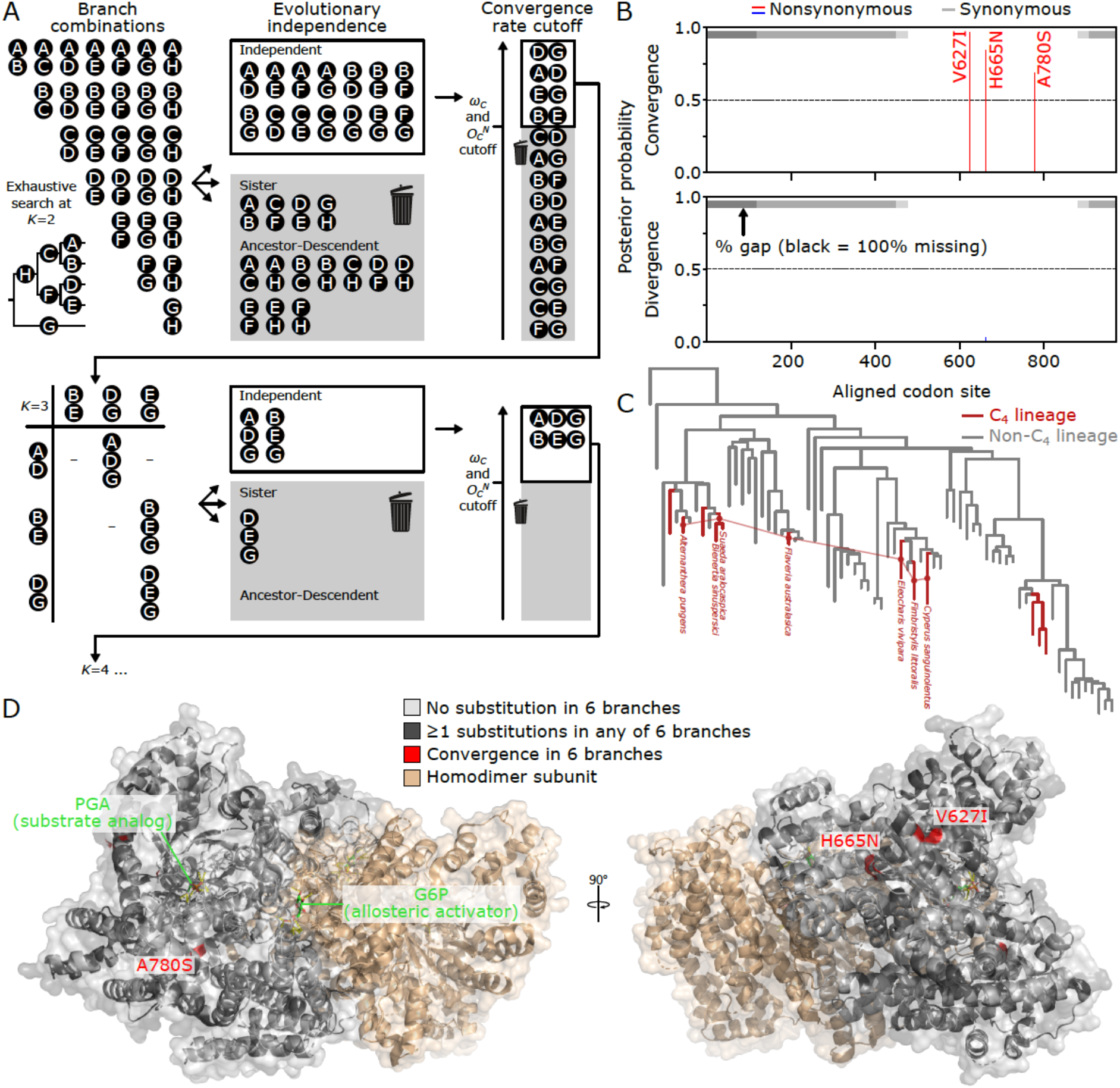
Analysis of highly repetitive convergence. (**A**) Overview of the new branch-and-bound algorithm. This is a detailed illustration of Fig. 4A. (**B**) Site-specific probabilities of combinatorial substitutions in PEPC at *K* = 6. (**C**) Convergent branch combination in the PEPC tree at *K* = 6. (**D**) Positions of higher-order convergent substitutions in the structure of maize PEPC (PDB ID: 6MGI) (Muñoz-Clares et al., 2020). Abbreviations: PGA, phosphoglycolate (substrate analog); G6P, glucose-6-phosphate (allosteric activator).

**Figure S11.**
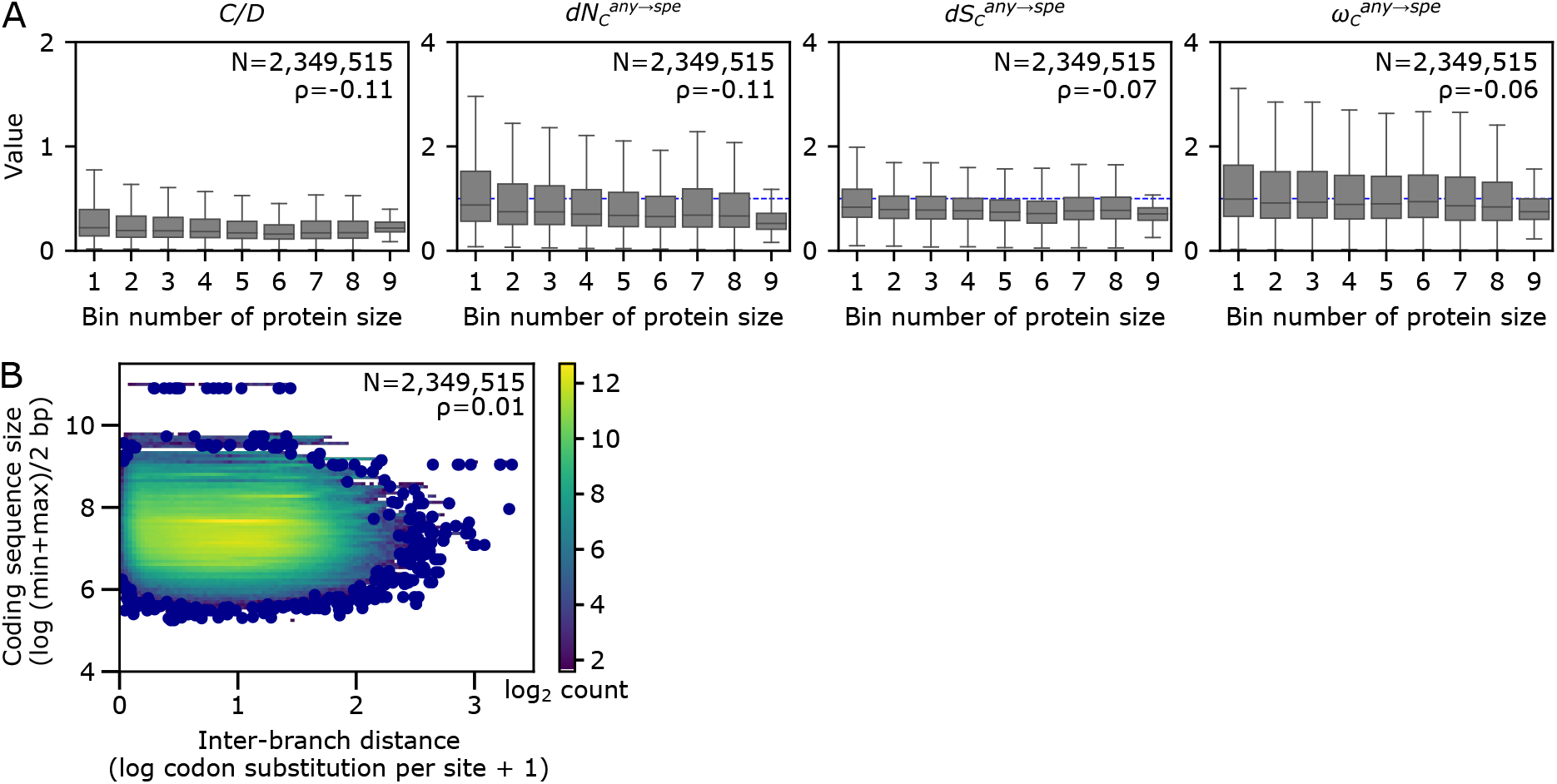
Relationships between protein sizes and convergence rates in vertebrate nucleus-encoded genes. (**A**) Protein-size-dependent variation of convergence rates. (**B**) Relationships between genetic distance and the size of proteins. While the inter-branch distance was obtained for each branch pair, the coding sequence size was defined for each orthogroup.

**Figure S12.**
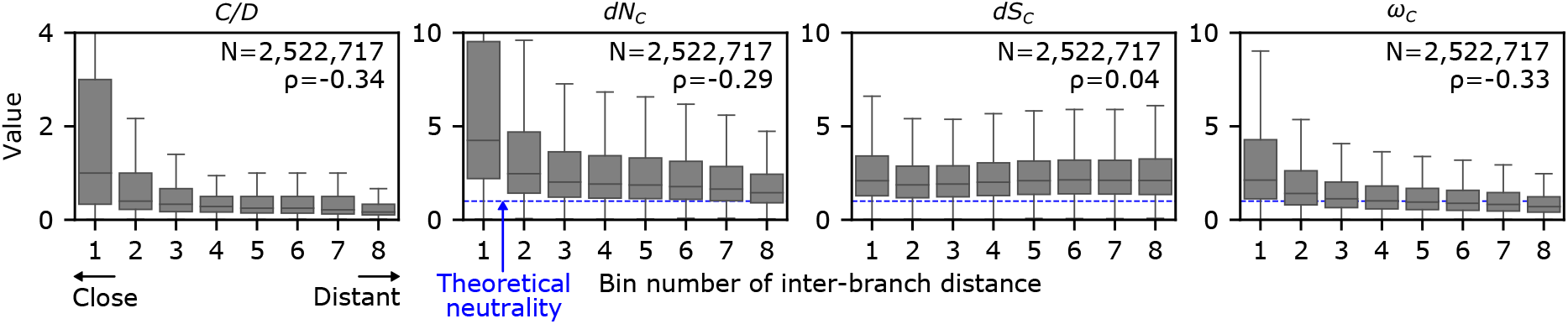
Temporal variation of convergence rates, as estimated with the binarized probabilities of ancestral states. The analysis of Fig. 2B is reproduced with the --ml_anc option in CSUBST. The number of branch pairs (N) and Spearman’s correlation coefficients (ρ) are provided in each plot. The bin range was determined to assign an equal number of branch pairs. To reduce the noise originating from branches where almost no substitutions occurred, branch pairs with both 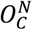 and 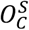 greater than or equal to 1.0 were analyzed.

**Figure S13.**
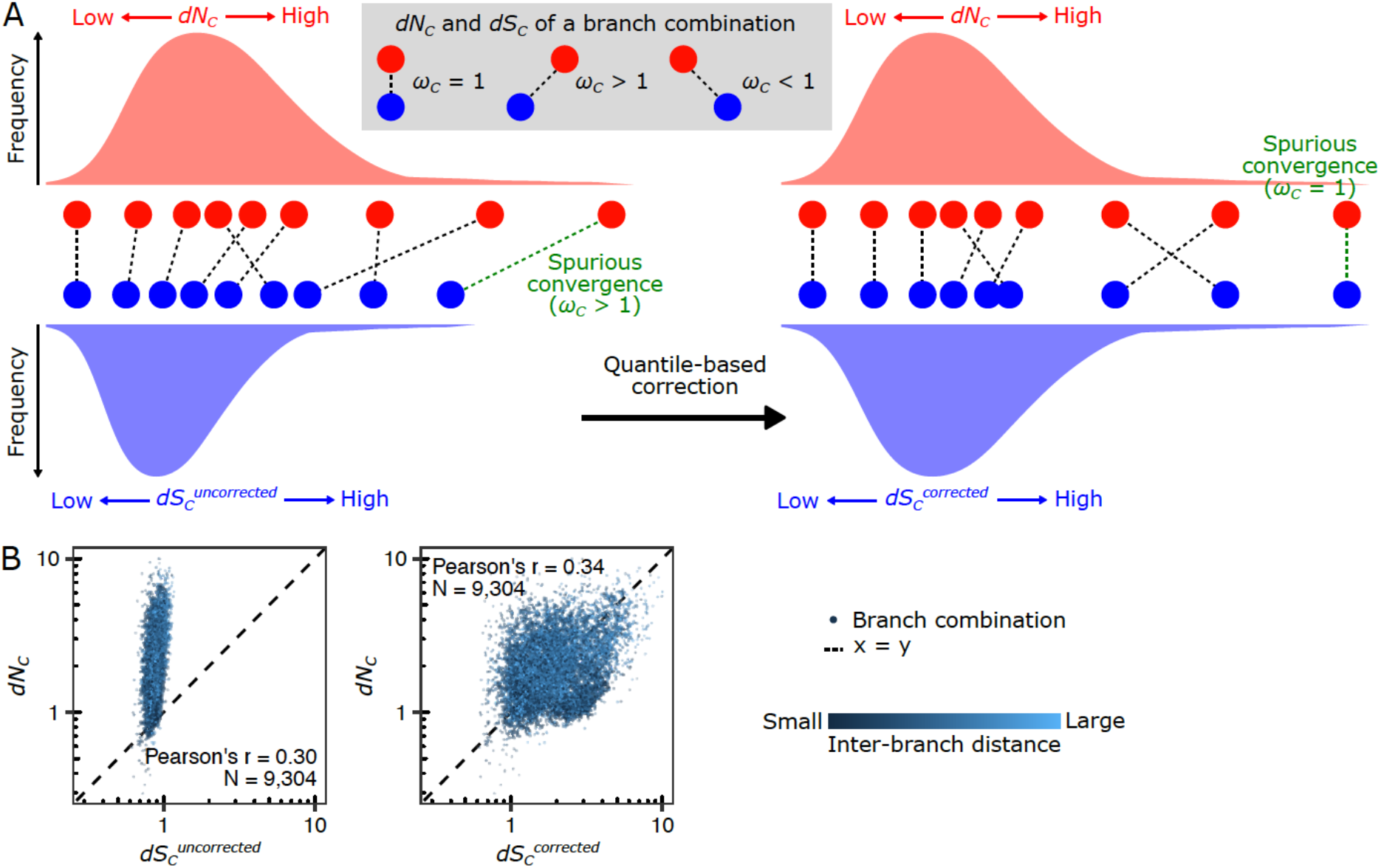
The long-tail correction matches the range of distributions between *dN_C_* and *dS_C_*. (**A**) A schematic representation of the long-tail correction (Equation 18). (**B**) Calibration of synonymous convergence rates in mitochondrial proteins. The mitochondrial genome data in Fig. 1E was analyzed. The inter-branch distance is shown on a color scale. The number of branch pairs (N) and Pearson’s correlation coefficients (r) are provided in the plot.

